# Out of sight: Surveillance strategies for emerging vectored plant pathogens

**DOI:** 10.1101/2022.01.21.477248

**Authors:** Alexander J. Mastin, Frank van den Bosch, Yoann Bourhis, Stephen Parnell

## Abstract

Emerging pests and pathogens of plants are a major threat to natural and managed ecosystems worldwide. Whilst it is well accepted that surveillance activities are key to both the early detection of new incursions and the ability to identify pest-free areas, the performance of these activities must be evaluated to ensure they are fit for purpose. This requires consideration of the number of potential hosts inspected or tested as well as the epidemiology of the pathogen and the detection method used. In the case of plant pathogens, one particular concern is whether the visual inspection of plant hosts for signs of disease is able to detect the presence of these pathogens at low prevalences, given that it takes time for these symptoms to develop. One such pathogen is the ST53 strain of the vector-borne bacterial pathogen *Xylella fastidiosa* in olive hosts, which was first identified in southern Italy in 2013. Additionally, *X. fastidiosa* ST53 in olive has a rapid rate of spread, which could also have important implications for surveillance. In the current study, we evaluate how well visual surveillance would be expected to perform for this pathogen and investigate whether molecular testing of either tree hosts or insect vectors offer feasible alternatives. Our results identify the main constraints to each of these strategies and can be used to inform and improve both current and future surveillance activities.

## Introduction

Increases in international travel, transportation, and trade have increased the risk of introduction of plant pests and pathogens into new areas, with changes in land use and climate potentially facilitating their establishment and spread (Anderson et al., 2004; Brasier, 2008; Waage and Mumford, 2008). Surveillance activities in presumed “pest free areas” (IPPC, 2021) are required to either confidently declare pest or pathogen absence (in order to facilitate trade activities) or to detect new incursions at a sufficiently early stage for control measures to be applied (Parnell et al., 2017), and are thus commonly referred to as “detection surveys”. To date, detection surveys are generally based upon the visual inspection of economically or ecologically important host species by trained surveyors (IPPC, 2021). Whilst this strategy is invaluable for the detection of novel and unexpected pests and pathogens, there are concerns that it may be less effective in cases where the pest or pathogen is known but symptoms do not immediately become apparent. This is evidenced by the fact that many emerging pests and pathogens are first detected at a point in epidemic development at which control is no longer feasible (Cunniffe et al., 2016; Gottwald et al., 2006; Herms et al., 2004; Sansford, 2013). Our own previous work has demonstrated that, along with the epidemiology of the pathogen, the detection method has an effect on the number of hosts which must be inspected or tested for detection surveys to be effective (Alonso Chavez et al., 2016; Bourhis et al., 2019a; Mastin et al., 2017, 2019; Parnell et al., 2012, 2015). Although new diagnostic methods capable of detection of infection in presymptomatically infected hosts (Silva et al., 2021) offer great potential for improving detection in individual hosts, less is known about their value for large scale detection surveys – particularly as they will generally cost more than visual inspection to deploy (Mastin et al., 2020). In the case of molecular tests, there is also the question of whether hosts should be tested at all. Many plant viruses and some notable bacterial plant pathogens are spread by insect vectors, which may themselves be valuable alternative sources of surveillance data (Mastin et al., 2017), yet are generally only used as an adjunct to conventional host-based surveillance.

The challenges facing visual inspection as a surveillance strategy are exemplified by the recent emergence of a novel strain of the vector-borne plant pathogenic bacterium *Xylella fastidiosa* in Europe. This strain (*X. fastidiosa* subspecies pauca, ST53 – hereafter *X. fastidiosa* ST53) was identified in 2013 as the cause of a novel disease of olive trees (olive quick decline syndrome; OQDS) in the Italian province of Lecce in the region of Apulia (Martelli et al., 2015; Saponari et al., 2013). Following first identification, the meadow spittlebug, *Philaenus spumarius*, was identified as the most important vector of this pathogen (Ben Moussa et al., 2016; Cornara et al., 2016, 2017a; Saponari et al., 2014) and the limits of infection within the Salento peninsula were identified through a delimiting survey. Whilst elimination of infection from this area is considered unlikely, *X. fastidiosa* is considered one of the greatest phytosanitary threats in Europe, meaning that there is now a need for effective surveillance in areas considered still free of infection (European Commission, 2021). Although much of this surveillance to date has been based upon visual inspection, the long “presymptomatic period” before hosts become visibly detectable and the high potential spread rates of *X. fastidiosa* raise questions of the efficacy of this strategy (EFSA et al., 2020a). By building upon our earlier work (Bourhis et al., 2019a; Mastin et al., 2017, 2019; Parnell et al., 2015), we consider here whether visual detection can continue to be justified as the standard surveillance strategy prior to *X. fastidiosa* incursion, in the face of alternatives such as molecular testing of either hosts or vectors by considering the following questions:

- Is visual inspection useful for detection surveys?
- What characteristics of a host diagnostic test would make it more cost effective than visual inspection?
- Could laboratory testing of vectors outperform visual inspection?

## Methods

### Is visual inspection useful for detection surveys?

A single detection survey can result in one of two potential outcomes:

i. At least one positive detection is made, usually after a series of monitoring rounds where no detections are made. Assuming that the detection method in use has a perfect specificity – that is, there are no ‘false positive’ results – this indicates that the pathogen is definitely present in the population.
ii. No positive detections are made, in which case the pathogen may or may not be present in the population, due to imperfect test sensitivity (i.e. ‘false negative’ results) and/or random error (i.e. the possibility that infected hosts are present but were not sampled).

Whether or not the pathogen of interest is found during this detection survey, we are interested in answering the same general question: “given these results, what can we say about the prevalence of infection in this area?”. If a pathogen is not detected, we commonly reformulate this question in relation to a predefined “prevalence threshold” and ask what the probability is that the prevalence is lower than this threshold. If this probability is sufficiently high, it can be interpreted as evidence that the pathogen is effectively absent from the region in question (EFSA et al., 2020b; European Commission, 2021). This interpretation links well with our previous work on pest freedom determination (Bourhis et al., 2019a), which allows us to estimate the probability density, *P*(*q*), of the prevalence, *q*, for any given sampling rate and thus identify the prevalence above which only a small percentage of the probability density remains. By changing the number of hosts inspected (and found to be negative), we can estimate the number of hosts which would need to be sampled in order for this prevalence to be lower than a given prevalence threshold, and therefore confidently declare pest freedom. For ease of calculation, we consider a single detection survey, allowing us to disregard the interval between survey rounds (Bourhis et al., 2019a). However, our methods are also applicable to multiple rounds of a detection survey across years (either to determine pest freedom or in the case of early detection), as detailed in Supplementary Information A.

When our detection survey is based upon visual inspection, our ability to detect infection will depend upon the proportion of symptomatic hosts at any given time, which we term the “apparent prevalence”. However, we wish to declare pest freedom in relation to the true prevalence (the proportion of infected hosts, symptomatic or not). The relationship between the apparent and true prevalences will be affected by both the duration of the presymptomatic period (which we term the “detection lag”) and the rate of pathogen spread (*r*) (Figure 1). The ratio of true and apparent prevalences would also be expected to reduce over time as density dependent constraints reduce the rate of increase in the true prevalence, until the true and apparent prevalences are equal (Figure 1B). As this effect will be most pronounced when a pathogen is spreading rapidly and the detection lag is relatively long (as is the case with *X. fastidiosa* ST53), we explicitly consider this logistic growth pattern in our model (Bourhis et al., 2019b), rather than the exponential approximation (i.e. an assumed fixed ratio of true and apparent prevalences over time) we have previously described (Alonso Chavez et al., 2016; Bourhis et al., 2019a). Using this approach, we are able to estimate the number of trees that would need to be visually inspected to detect a maximum true prevalence of 0.01 at a confidence level of 0.90 (Supplementary Information A), for a range of different tree pathogens – including *X. fastidiosa*.

**Figure 1.**
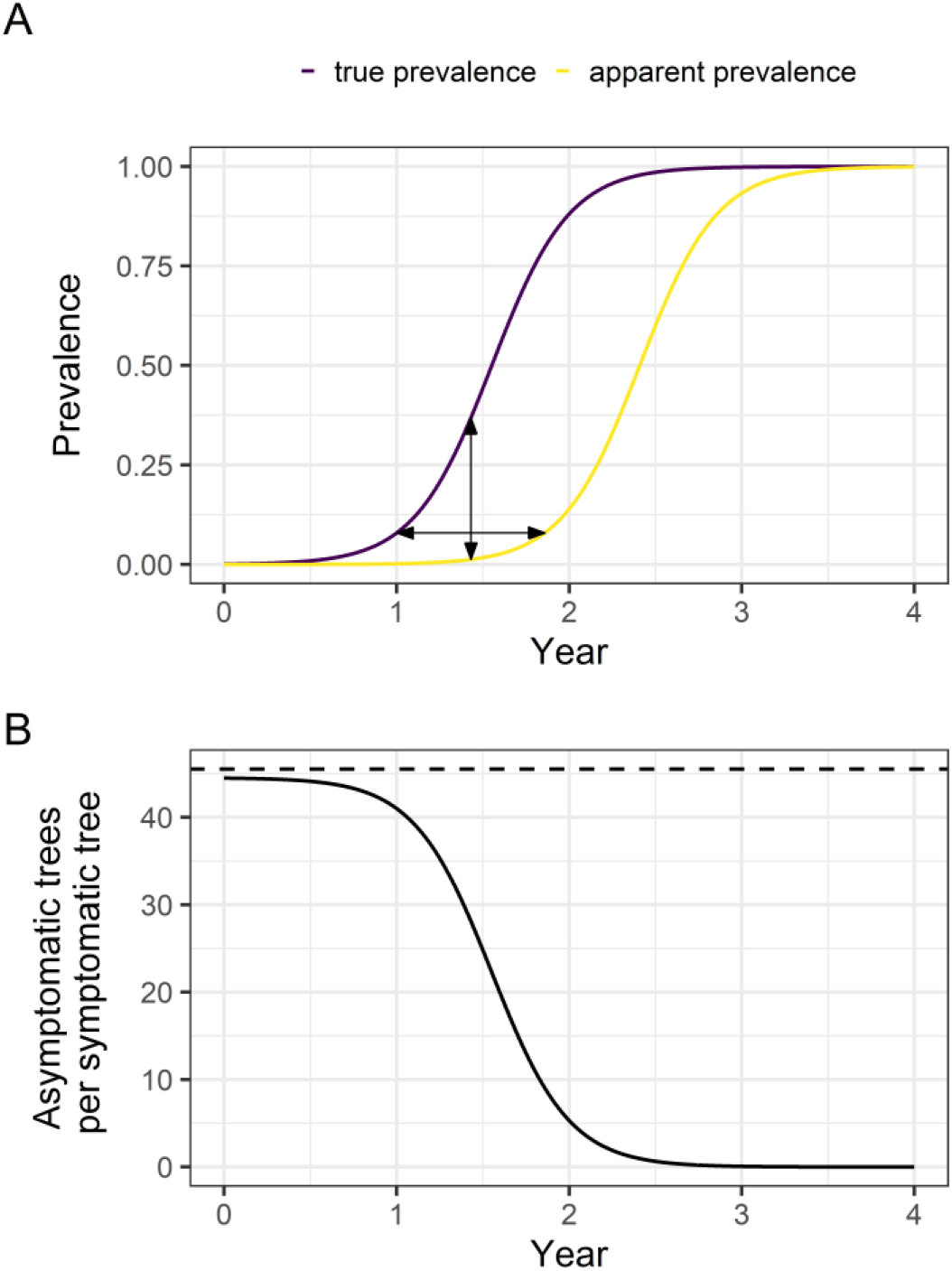
The true prevalence of infection can be estimated for any apparent prevalence from the rate of pathogen spread and the length of the detection lag period. **A. A detection lag period can be considered as a shift of the epidemic growth curve to the right.** In this plot, time is shown on the x-axis and the proportion of infected or symptomatic hosts (the prevalence) on the y-axis. The two curves represent the true prevalence and the apparent prevalence (e.g. the proportion of hosts with symptoms, if detection is based upon visual inspection). The horizontal distance between the curves (i.e. in the direction of the x-axis) represents the asymptomatic period (the detection lag (*δ*) for visual inspection), and the vertical distance (in the direction of the y-axis) represents the difference between the true and apparent prevalences at any given time. **B. The ratio of the true and apparent prevalences decreases as the true prevalence increases.** This plot shows the ratio of the true and apparent (detectable) prevalences (which can be interpreted as the number of asymptomatic trees per symptomatic tree) under logistic and exponential growth as time progresses. The dashed line represents the predicted ratio under continued exponential growth and the solid line represents that under logistic growth. Although during very early stage spread, the growth in both the true and apparent prevalences is broadly exponential, as the true prevalence deviates from this, the ratio of the two prevalences starts to decrease.

### What characteristics of a host diagnostic test would make it more cost effective than visual inspection?

Can we improve upon the performance of a detection survey by using a laboratory test capable of identifying infection in presymptomatic hosts? The methods described above allow us to explore the impact of varying different test characteristics (namely the detection lag period and diagnostic sensitivity), but we also need to consider how the costs of alternative detection methods compare to those of visual inspection. To do this, we adapt our previous work on early detection surveillance (in which the pathogen is detected) (Mastin et al., 2019) to the situation in which there is no detection (Supplementary Information A). Rather than specifying a specific detection method, we investigate what combinations of detection lag and diagnostic sensitivity and relative cost would be required to outperform visual inspection, assuming a single round of sampling. However, we also consider the specific example of a molecular diagnostic which costs €14.63/host to deploy, in contrast to visual inspection at €5.48/host, based on estimates of the costs of *X. fastidiosa* surveillance in Apulia (Supplementary Table 2).

### Could laboratory testing of vectors outperform visual inspection?

Vector-borne pathogens such as *X. fastidiosa* can be detected in insect vectors as well as in the plant host. Because *X. fastidiosa* is a ‘semipersistent’ pathogen (Almeida et al., 2005; Purcell and Finlay, 1979), it is restricted to the foregut of infected vectors, meaning that colonised tissue can be more reliably isolated in infected insect vectors than in infected plant hosts. However, the question of what proportion of vectors are infected, and how this compares to that in plant hosts, remains. Not only is very little known about how the prevalence of *X. fastidiosa* in vectors and hosts relate to each other during early stage spread, there is also marked seasonality in vector infection. *P. spumarius* is univoltine (i.e. a single new generation is produced per year) and adults rarely survive the winter months. The total density of adult vectors therefore rapidly increases from the time of first emergence in spring, to peak in summer, before decreasing to very low levels over the winter months due to mortality (Cornara et al., 2017b). As *X. fastidiosa* is lost during moulting and is not transmitted vertically, adult vectors (which are motile and therefore the main source of tree to tree spread (Cornara et al., 2018)) would be expected to only acquire infection in a relatively short window following emergence in spring and whilst feeding on potentially infected olive hosts (before moving to herbage in late summer). In Apulia, the prevalence of vector infection therefore reduces to very low levels over winter each year before increasing in the Spring and Summer months (Ben Moussa et al., 2016; Cornara et al., 2017b, 2017a).

We capture the temporal trends in both *P. spumarius* and olive infection using an epidemiological model (described in more detail in Supplementary Information B), allowing us to estimate how the density of P. spumarius varies over the course of a year (Figure 2A), and how the prevalence of *X. fastidiosa* in vectors and hosts changes both within and between seasons (Figure 2B). We parameterise this model using data on both the abundance of adult *P.spumarius* (Ben Moussa et al., 2016; Cornara et al., 2017b) and their prevalence of infection with *X. fastidiosa* (Ben Moussa et al., 2016; Cornara et al., 2017b, 2017a) over the course of a year. We captured the associated prevalence of infection in olive hosts by estimating the overall mean prevalence in these hosts over the same time period (between 2013 and 2015) and in the same area of Lecce province captured in the vector data. At the same time the peak vector prevalence was 0.48 (Figure 2B), the prevalence amongst hosts was 0.23. We then simulate early-stage spread between vectors (accounting for the seasonal trends in both density and prevalence) and hosts (in which the prevalence increases over consecutive years according to the total density of bacteria-carrying vector days over the course of the previous year). Model predictions of host and vector prevalence over five consecutive years are shown in Figure 3B.

**Figure 2.**
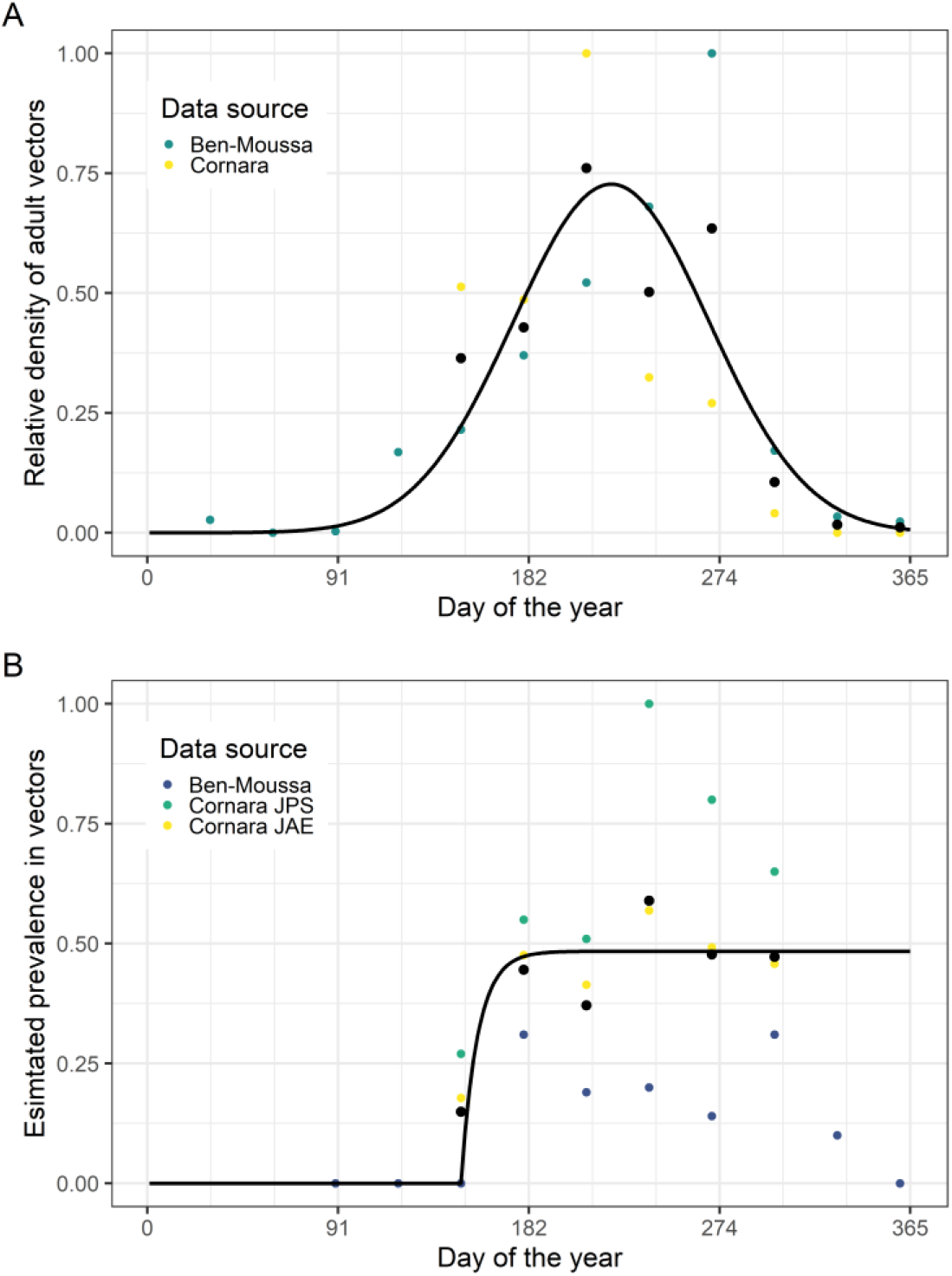
There is pronounced seasonal variability in the density of adult P. spumarius and the prevalence of X. fastidiosa infection amongst these. **A: Data suggest that adult *P. spumarius* are absent from January to March, and peak in density around August.** This plot shows the modelled change in relative *P. spumarius* density over a year, fitted to data from two papers. The black dots show the mean density from both papers. **B: Data suggest that the prevalence of *X. fastidiosa* infection in adult *P. spumarius* increases rapidly between June and July, to reach a steady peak for the rest of the year.** This plot shows the modelled change in the prevalence of *X. fastidiosa* infection of *P. spumarius* over a year, fitted to data from three papers. “Cornara JPS” is (Cornara et al., 2017b) whereas “Cornara JAE” is (Cornara et al., 2017a) The black dots show the mean prevalence estimates from all three papers.

**Figure 3.**
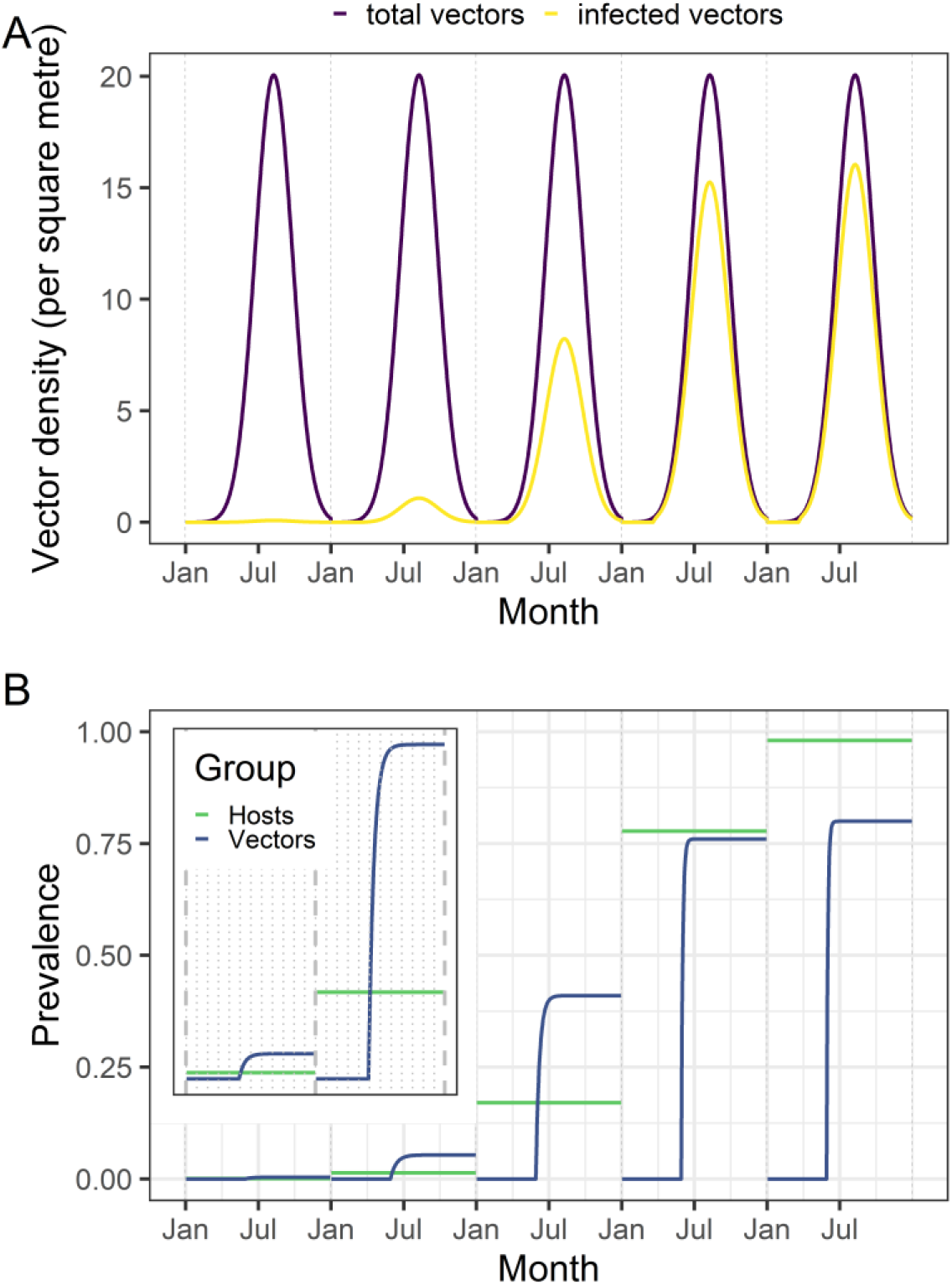
In the early stages of the epidemic, the apparent prevalence of X. fastidiosa in vectors increases faster than that in hosts. **A: Although the total density of *P. spumarius* is assumed to be fixed between years, the density of infected vectors increases each year.** This plot shows the modelled density of *P. spumarius* and the density of *X. fastidiosa*-infected *P. spumarius* over the course of five years. **B: The prevalence of *X. fastidiosa* infection is higher in vectors than in hosts in the early stages of a new epidemic.** This plot shows how the modelled prevalence of *X. fastidiosa* in hosts contrasts with that in vectors, over the course of five years. The inset plot shows the estimates from the first two years in more detail.

We assess the implications of these results for surveillance by adapting our previous work on early detection surveillance in a host-vector system (Mastin et al., 2017) for a scenario in which no detections are made (as described in Supplementary Information C). This approach requires a single estimate of the ratio of apparent vector and host prevalences during early stage spread. As this ratio varies both within and between years in the case of *X. fastidiosa* (Figure 3B), we consider only very early stage spread. If we assume that vector sampling is conducted when vector densities (and the prevalence of infection) is at its peak (which is most logistically feasible and therefore commonly practiced in the field), we can estimate the initial ratio of detectable vectors and detectable hosts either analytically (see Supplementary Information C for more information) or directly from the model. Using this estimate, we can then estimate the maximum apparent prevalence in hosts for any given number of hosts and/or vectors inspected/tested and found to be negative, and convert this to an estimate of the true host prevalence under the assumption of logistic growth using the methods described above. As our previous work has shown that the total surveillance costs required in order to detect infection at or before a given prevalence are generally minimised when either hosts only or vectors only are sampled (Mastin et al., 2017), we only consider these two scenarios here (rather than a mixed surveillance strategy in which both hosts and vectors are sampled). Finally, we incorporate sampling and testing costs and estimate the total costs of either host or vector sampling. As the relatively low numbers of bacteria in infected individuals (Cornara et al., 2016; Hill and Purcell, 1995a, 1995b) limit the ability to detect *X. fastidiosa* infection in vectors using ELISA tests (Huang et al., 2006), we consider PCR testing (Harper et al., 2010) of vectors here.

## Results

### Is visual inspection useful for detection surveys?

The number of hosts which must be found to be asymptomatic to declare pest freedom is affected by the rate of pathogen spread and the duration of the presymptomatic period, which will vary for different tree pathogens. These factors together determine the degree of disparity between the apparent prevalence (the proportion of hosts with visual symptoms) and the true prevalence of infection. Using the mean estimates of spread rate and presymptomatic period summarised in (Alonso Chavez et al., 2016), we find that there is little difference between the apparent and true prevalences for some pathogens (such as *Phytophthora ramorum* or *Hymenoscyphus fraxineus*). In these cases, relatively small numbers of hosts must be inspected in order to be able to declare pathogen freedom (Figure 4). However, using our own estimates for *X. fastidiosa* ST53 (Supplementary Table 2), we found that the disparity between the apparent and true prevalences was more marked than for any other pathogen considered, with around 80% of hosts being infected by the time 10% become detectable (Figure 4A). The low numbers of symptomatic hosts which would be expected during early stage spread means a total of 10,384 trees would need to be observed (and all found to be asymptomatic) to be 90% confident that the prevalence of *X. fastidiosa* ST53 was lower than 1%, under our best estimates of the growth rate (0.0122 infections/infected host/day) and presymptomatic period (313 days). This is greater than for any of the other pathogens considered (Figure 4B).

**Figure 4.**
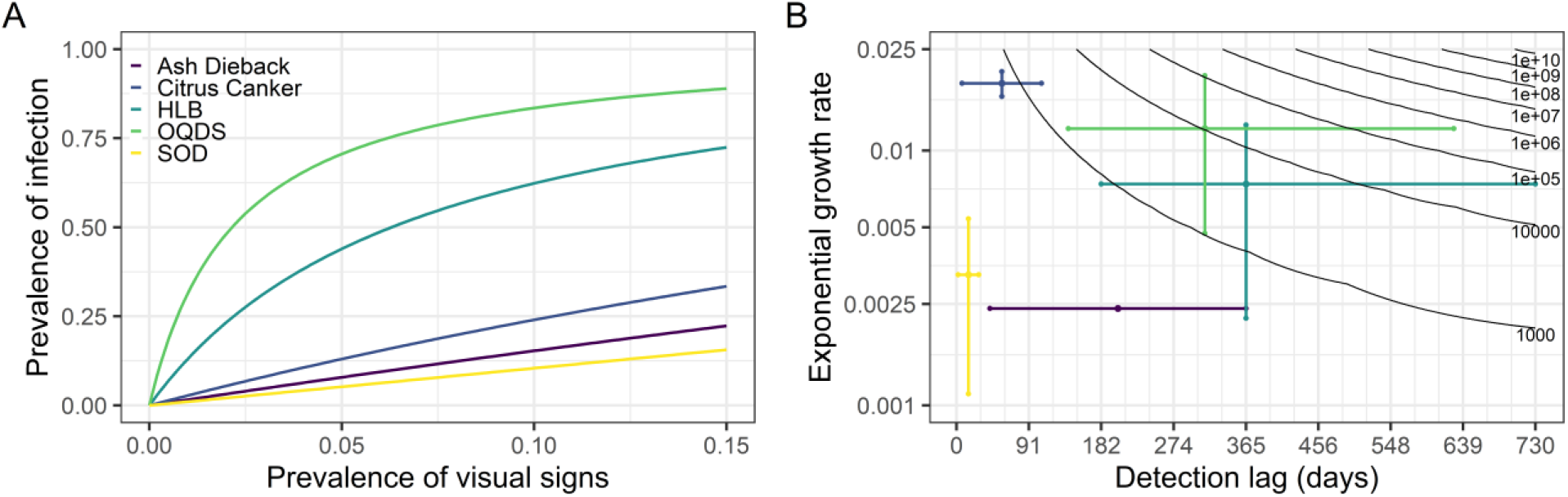
The asymptomatic period for *Xylella fastidiosa* makes it very difficult to detect at an early stage when using visual inspection. **A: The highest difference between apparent and true prevalence is seen for olive quick decline syndrome, caused by *X. fastidiosa*.** This plot shows the relationship between the apparent prevalence (on the x-axis) and the true prevalence (on the y-axis) for a number of different diseases (causal pathogens): ash dieback (*Hymenoscyphus fraxineus);* citrus canker (*Xanthomonas axonopodis* pv. *citri);* huanglongbing/HLB (*Candidatus* Liberbacter asiaticus); olive quick decline syndrome/OQDS (*X. fastidiosa* ST53); sudden oak death (*Phytophthora ramorum*). **B: In order to confidently declare pest freedom, more samples are needed when the asymptomatic period and/or the spread rate are high.** This plot shows the relationship between the detection lag (x-axis), the exponential growth rate (on a logarithmic scale on the y-axis), and the number of samples (also on a logarithmic scale, in the contour lines) required to be 90% confident that the true prevalence is lower than 1% given that no positive detections are made. The coloured lines indicate our best estimates of the growth rate and presymptomatic period for the pathogens considered (Alonso Chavez et al., 2016).

### What characteristics of a host diagnostic test would make it more cost effective than visual inspection?

Although detection methods able to detect the pathogen before the development of symptoms (i.e. with shorter detection lags) require fewer samples to be collected (Figure 5A), any associated reductions in the diagnostic sensitivity increases the required sample size. There is therefore a tradeoff between the detection lag and the diagnostic sensitivity, meaning that both of these test characteristics must be considered together. Although the most marked reduction in sample size is associated with relatively small reductions from the original detection lag, there are considerable increases in required sample size when the sensitivity of detection is low (Figure 5A). In practical terms, a test with a detection lag period half that of visual inspection would require fewer samples than visual inspection if the diagnostic sensitivity of this test was over 0.15.

**Figure 5.**
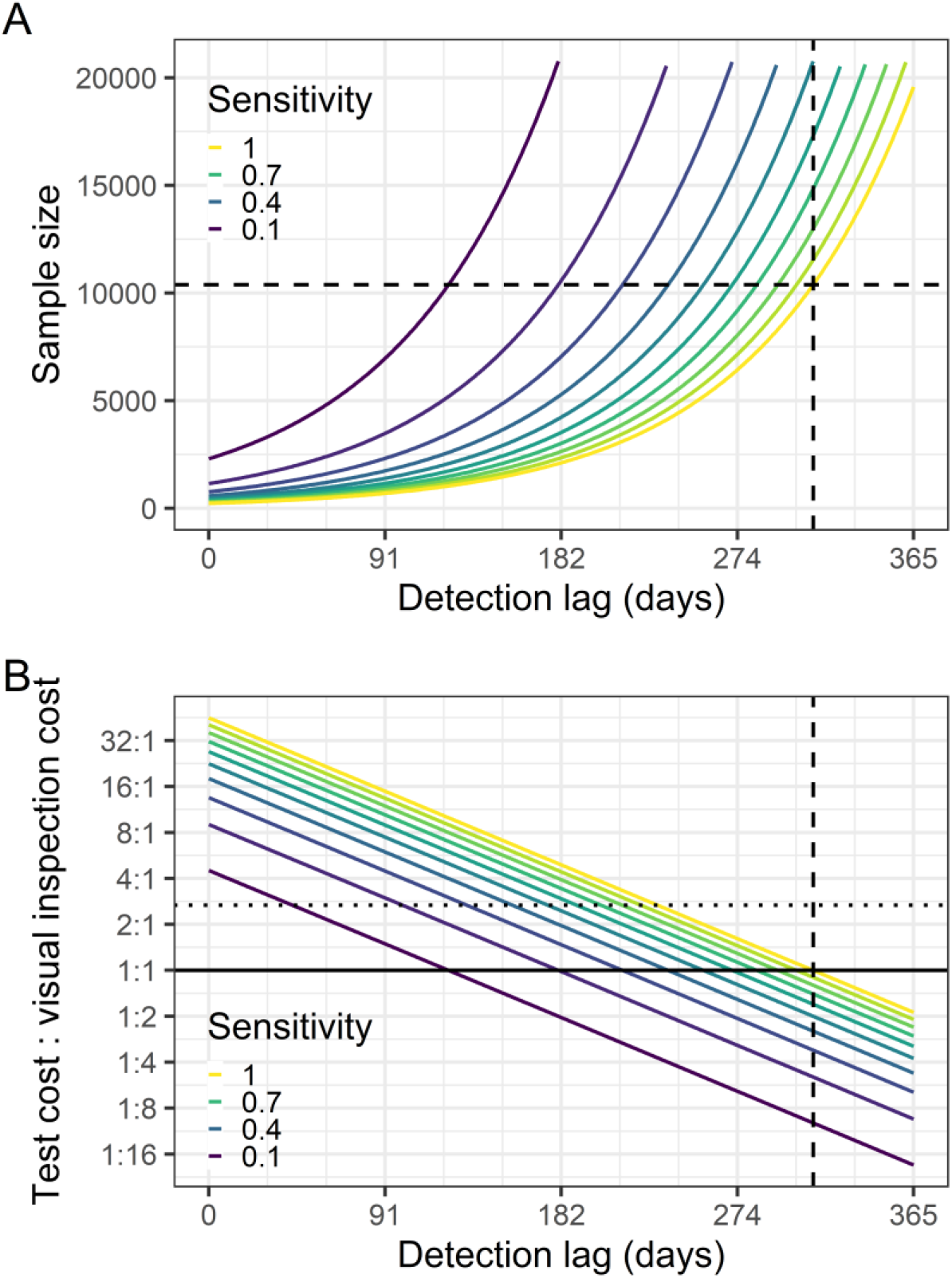
The low sensitivity of current diagnostic tests when applied to presymptomatic hosts may limit their ability to detect infections at a low prevalence. **A: Diagnostic tests can result in a lower sample size than visual inspection, but if the diagnostic sensitivity is low, the test needs to be able to detect infection shortly after infection.** This plot shows the impact of reducing the detection lag and the diagnostic sensitivity on the number of hosts which must be found to be negative to be 90% confident that the prevalence is lower than 1% (the sample size). As the dashed lines reflect the detection lag and required sample size under visual inspection, all solid lines below the horizontal dashed line indicate that fewer trees must be tested to declare pathogen freedom than would have to be visually inspected. **B: Lower detection sensitivities and higher costs both reduce the feasibility of a nonvisual detection method, even if the detection lag is short.** This plot expands on plot A to also incorporate testing costs. We capture this by showing the in solid coloured lines the relative cost of an alternative detection method at which the total costs of surveillance would be equal to those under visual inspection, for different diagnostic sensitivities. The intersections of the solid coloured lines with the solid black line (which indicates that the costs of the detection method is equal to that of visual inspection) therefore represent equal required sample sizes (and therefore match the intersections of the curves in plot A with the dashed line). The horizontal dotted line indicates the current estimated relative cost of using the host ELISA test (a cost ratio of €14.63/€5.48 =2.67). The vertical dashed line shows the presymptomatic period for *X. fastidiosa* (and therefore the detection lag for visual inspection). All areas of the parameter space below the test sensitivity contour of interest indicate that the alternative detection method is cheaper to deploy than visual inspection, and all areas above indicate that visual inspection is cheaper.

However, the detection lag and diagnostic sensitivity are not the only important considerations for an alternative detection method. We also need to consider how much the new method costs to deploy, and how this compares to visual inspection (Mastin et al., 2019). Assuming that the alternative detection method is an ELISA test, which costs around 2.67 times more than visual inspection to deploy (Maria Saponari, Personal Communication), we find that a test with a perfect sensitivity must be able to detect infection with *X. fastidiosa* at or before 232 days post-infection to be more cost effective than visual inspection (as shown in the horizontal dotted line in Figure 5B). If the sensitivity of the test is lower, then it must be possible to detect infection even earlier than this for the test to be more cost effective than visual inspection. Although very little information is available on the performance of the ELISA test on asymptomatically infected hosts at different times post-infection, it is likely to be very low, given the large number of leaves (the majority of which will not contain bacteria) on a tree.

Our method can also be used to assess other potential host-based detection methods. Figure 5B shows how the detection lag, diagnostic sensitivity and cost influence the required sample size together by estimating the “equivalence point” at which the total cost of either visual inspection or the alternative detection method would be equal for any combination of these three factors. This equivalence point is shown in Figure 5B for different combinations of diagnostic sensitivity, detection lag, and relative test cost (the ratio of the costs of testing a single host with the alternative detection method and by visual inspection) as coloured lines. For any given diagnostic sensitivity (i.e. selecting a single coloured line in Figure 5B), we find that changing the relative costs effectively shifts the previous relationship between detection lag and required sample size in a linear fashion. This means that doubling the relative costs of the molecular test reduces the maximum acceptable detection lag by 57 days, all else being equal.

### Could laboratory testing of vectors outperform visual inspection?

To evaluate how vector testing would be expected to compare to host visual inspection we need to consider not just the diagnostic considerations of detection lag, diagnostic sensitivity, and cost, but also any differences in the prevalence of infection between hosts and vectors, which will be determined by the epidemiology of the pathogen itself. We find that these epidemiological considerations are favourable for vector surveillance during early stage spread, with the prevalence of vector infection being up to four times higher than that in hosts. Although the detection lag and diagnostic sensitivity of a PCR test are also favourable for vector surveillance, the higher costs associated with such testing means that vectors must be pooled in order for these approaches to be cost-effective.

Our model of the population dynamics of adult *P. spumarius* replicates the seasonal fluctuations in adult *P. spumarius* density (Figure 3A) and prevalence of infection with *X. fastidiosa* (Figure 6A) seen in the data (Figure 2). In line with the available data (Figure 2B), the prevalence of infection is zero when adults are not present, before rising as adults emerge and initially feed on olive hosts (EFSA et al., 2019; Fierro et al., 2019), and then remaining unchanged for the remainder of the year as the total density of adults declines (which we term the asymptotic prevalence). Our model is also able to predict how the prevalence of *X. fastidiosa* amongst adult *P. spumarius* varies over a number of years (Figure 3B). Despite the similar general trend each year, the asymptotic prevalence in vectors increases over the first four years, as does the prevalence in hosts (Figure 3B). However, these are not symmetrical increases – with the vector prevalence reducing from 4.06 times higher than the host prevalence in the first year to 3.86 times higher in the second year and 2.40 times higher in the third year. The fact that this ratio remains greater than 1.0 shows that during early stage spread, any given number of sampled vectors would have a higher probability of containing an infected vector than an equal number of sampled hosts.

**Figure 6.**
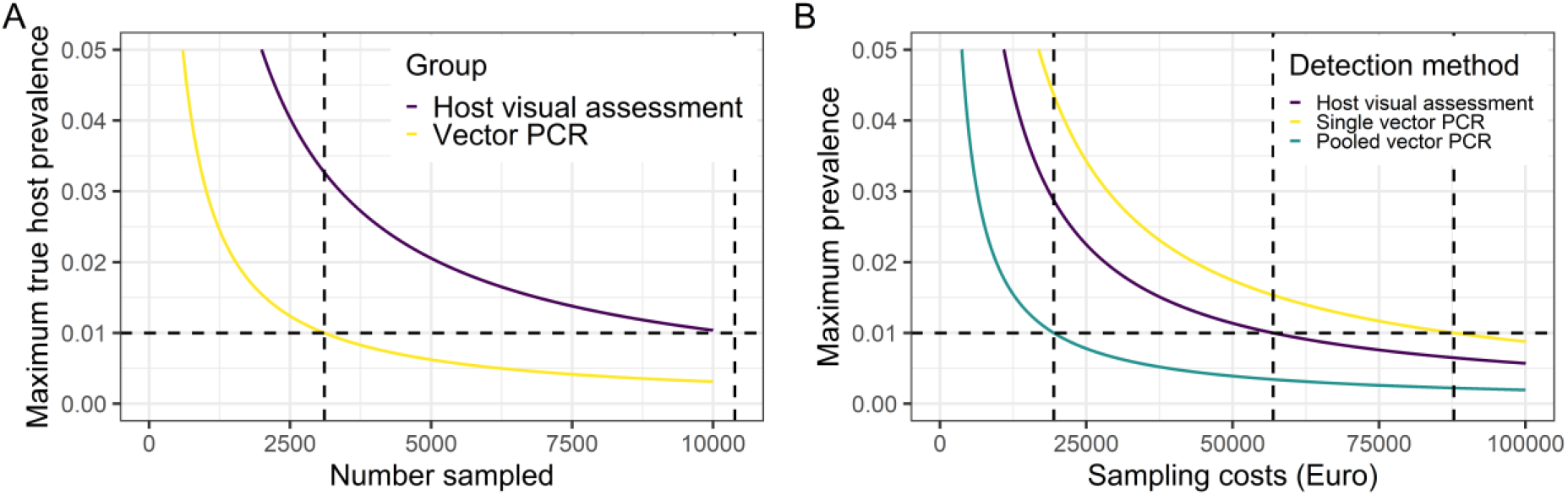
In cases where *X. fastidiosa* is not thought to be present, fewer vectors than hosts need to be tested in order to declare pest freedom. **A: The number of hosts which need to be tested to detect at a given prevalence is over three times higher than the number of vectors.** This plot shows the required number of individuals to be sampled on the x-axis and the 95^th^ percentile of the host prevalence in the absence of positive detections on the y-axis, when hosts (red) or vectors (blue) are sampled exclusively. The intersection of the curves and the horizontal dashed line represents the sample size required to be 90% confident that the true prevalence is lower 1% if no detections are made. **B: If vectors are pooled, the total cost of sampling hosts is around three times higher than the cost of sampling vectors.** This plot shows the total sampling costs on the x-axis and the 95^th^ percentile of the host prevalence in the absence of positive detections on the y-axis, when hosts (red) or vectors (blue and green) are sampled exclusively. We assume that hosts are sampled with visual inspection and ELISA confirmation of suspected positives, and that vectors are tested using qPCR, either singly (blue) or pooled in batches of five (green).

When we consider only the differences in detection lag and diagnostic sensitivity between vector and host sampling, we find that a total of 10,384 hosts would need to be sampled to be able to declare a prevalence lower than 1%, in contrast to 3,106 vectors (Figure 6A). However, when we account for the fact that laboratory testing of single vectors is higher than the costs of host visual inspection, we find that it would cost €87,754 to reliably declare pest freedom when sampling vectors in contrast to the €56,902 required for host visual inspection (Figure 6B). Studies have suggested that vectors can be pooled in batches of up to five insects (EPPO, 2019). We estimate that doing this would reduce the costs to €19,444, assuming that this pooling does not impact upon the test sensitivity (Figure 6B).

## Discussion

### Summary

The rate of new plant pathogen invasions has skyrocketed in recent years, associated with increases in international travel and trade and changes in land use and climate (Anderson et al., 2004; Brasier, 2008; Waage and Mumford, 2008). This is exemplified by the recent detection of the vector-borne plant pathogen *Xylella fastidiosa* in numerous European countries, reflecting a number of separate incursions (Landa et al., 2019). Given the considerable threats this pathogen poses to plant health throughout the continent were it to spread further, as well as the continued threat of incursion of other plant pathogens, we are faced with the question of how best to conduct surveillance to ensure that “pest free areas” remain as such. These surveillance activities (known as “detection surveys”) must be capable of detecting the presence of the pathogen at low prevalences, and to date, have predominantly relied on the visual inspection of host plants for signs of disease. Whilst this remains the only plausible method of detecting new, unexpected, pathogens or disease syndromes, it is unclear whether the wealth of alternative detection strategies offered by advances in molecular diagnostics and image analysis may be more appropriate when the pathogen of interest is known. We investigate whether visual surveillance can still be justified for *X. fastidiosa* detection surveys by comparing the expected performance of visual inspection to that for other tree pathogens and then evaluating the performance of alternative host-based methods such as molecular diagnostic tests and laboratory testing of insect vectors in relation to visual inspection. Although directly valuable for informing future surveillance for *X. fastidiosa*, our results allow us to better understand the situations in which these different detection methods may be best applied, and the constraints to their use.

Although *X. fastidiosa* has over 600 known potential host species (Castro et al., 2021), we focus here on the *X. fastidiosa* ST53 – olive system, as found in Apulia, Italy (Saponari et al., 2019). In this system, the combination of a high spread rate and a long presymptomatic period means that visual inspection is likely to fail to reliably detect invasions at an early stage of invasion (i.e. when the prevalence of infection is very low) unless very large numbers of hosts are inspected (Figure 4). Although fewer hosts would need to be inspected if molecular tests capable of reliably detecting infection before the development of symptoms were used, it is unlikely that this reliable detection can be achieved with these tests (since the probability of selecting a sample containing pathogen is so low). As a result of the likely low diagnostic sensitivity associated with molecular testing of presymptomatic trees, larger numbers of trees would have to be sampled (Figure 5A). As an additional constraint, the higher financial costs of molecular testing (even when using lower cost ELISA methods) compared to visual inspection also mean that sample size reductions would need to be very substantial before the tests become more cost effective (Figure 5B). However, there remains some promise in the use of higher throughput, whole-tree methods such as remote sensing, which may be capable of reliably detecting presymptomatic trees at a relatively low cost per tree (due to their capacity for inspecting large numbers of trees relatively quickly), which will be explored in more detail in future work. We also find that sampling insect vectors and testing them for the presence of the pathogen has the potential to outperform both visual inspection and molecular testing of hosts (Figure 6A). As well as offering shorter detection lag periods and higher diagnostic sensitivities, we find that the prevalence in vectors during early stage spread would be expected to be higher than that in hosts – making it more likely that infected individuals would be included in any sample, which therefore reduces the required sample size. The main challenge facing vector surveillance is the considerably higher per-sample costs of PCR testing. Although pooling vectors together for testing – a commonly used approach (EPPO, 2019) – may solve this problem (Figure 6B), further work is required to estimate the performance of this testing approach.

### Is visual inspection useful for detection surveys?

Visual inspection may be an appropriate detection method to use in pathogen detection surveys, but this depends on the rate of pathogen spread and the length of time before symptoms develop. As Figure 4 shows, the sample sizes required are lowest in cases where the pathogen spreads slowly and symptoms develop quickly (in Figure 4B, the lowest sample size contour is reached roughly when the time before symptoms develop is lower than the inverse of the spread rate). This agrees with our previous work, which showed that visual inspection for *Phytophthora ramorum* in rhododendron (a slow spreading pathogen with a short presymptomatic period) is likely to be more cost effective than the use of rapid diagnostic tests (Mastin et al., 2019). However, *X. fastidiosa* ST53 in olive both spreads rapidly and takes a long time for symptoms to develop. As a result, the number of trees which must be inspected for symptoms of *X. fastidiosa* infection during detection surveys is higher than for any other tree pathogen considered here (Figure 5B). Although this intensity of surveillance is comparable to that in recent years within the 10km wide ‘buffer zone (‘Zona Cuscinetto’) adjacent to the known infected zone in Apulia, it likely represents an unfeasibly high surveillance effort to maintain for long periods of time over the large areas for which such surveillance would be required (such as the remainder of Apulia, or even Italy as a whole). We assume that visual detection has both a perfect diagnostic specificity (i.e. that inspectors would be able to correctly identify all uninfected hosts) and a perfect diagnostic sensitivity (i.e. that inspectors would be able to detect all infected hosts after 313 days of infection). Our assumption of a perfect diagnostic specificity (i.e. that inspectors would not mistake other conditions for *X. fastidiosa* infection in uninfected hosts) corresponds to the guidance that any suspected cases would undergo confirmatory laboratory testing (European Commission, 2021), thereby making false positives unlikely. However, little is known of the true sensitivity of visual inspection for plant pathogens, due to variability in both symptom development and inspector performance. In the presence of a lower diagnostic sensitivity (for example, resulting from nonspecific or subtle symptom development), the required sample sizes would be further increased (Figure 5A).

### What characteristics of a host diagnostic test would make it more cost effective than visual inspection?

A number of novel methods of detection of host infection have become available in recent years. Although our method is flexible enough to be applicable to any of these, we focus mainly on the use of molecular tests – in particular, ELISA tests – which are currently being deployed in the field alongside visual inspection. Although theoretically capable of detecting presymptomatic infection in hosts and being relatively cheap to deploy, less is known of the diagnostic sensitivity of these tests in the field. Although estimates of test sensitivity are available, these are generally based upon the testing of either symptomatic or known infected tissue, and thus do not account for the fact that the pathogen is not homogeneously distributed throughout infected hosts – particularly early in infection. This is a particular issue for tree pathogens, and therefore means that there is a high probability that tissue sampled from an infected host will either not contain the pathogen at all or only at low levels. Given that even cheaper ELISA tests cost over two and a half times that of visual inspection, the number of hosts which need to undergo testing must be less than 40% of the number of hosts requiring visual inspection for the total surveillance costs of both methods to be equal. This can be achieved with a molecular test able to detect infection six months after infection (i.e. around five months before symptoms develop) and a host-level diagnostic sensitivity of greater than 0.7. To the authors’ knowledge, no available molecular tests could be expected to have sensitivities this high at this stage of infection. Although advances in molecular diagnostics offer some potential for reducing the costs of diagnostic tests, until a method of accurately selecting infected tissue for testing is developed, we therefore conclude that molecular testing of host plants is likely to remain of limited use for detection surveys.

Amongst nonvisual detection methods, non-molecular methods such as aerial remote sensing (Zarco-Tejada et al., 2018) or canine olfactory detection (Gottwald et al., 2020; Mendel et al., 2018) may be capable of more reliably identifying infected hosts before the development of symptoms as they operate at the level of the whole tree rather than a particular sample, and may therefore be more appropriate for detection surveys than either visual inspection or molecular testing. Another potential advantage of these particular methods is that they are potentially capable of screening large numbers of hosts in a short space of time, meaning that their cost of deployment on an individual host basis could be relatively low.

### Could laboratory testing of vectors outperform visual inspection?

We finally consider whether surveillance of insect vectors could circumvent some of the challenges associated with host surveillance. The concept of testing vectors for pathogens is a recognised component of surveillance for emerging vector-borne pathogens of humans and other animals (ECDC, 2012; Kading et al., 2018), as well as of plants (European Commission, 2021). Indeed, the first detection of the citrus pathogen *Candidatus* Liberibacter asiaticus (the cause of the citrus disease huanglongbing) in California was made in insect vectors (Kumagai et al., 2013). However, to date, most vector surveillance for *X. fastidiosa* has focused on the identification of competent vectors, seasonality of infection, and the spatial limits of the pathogen (Ben Moussa et al., 2017; Cruaud et al., 2018; Yaseen et al., 2015). We find that the high prevalences of vector infection during early stage spread, the potential for reliable detection early in infection, and the ability to reduce testing costs through pooling, all make insect vectors a valuable “sentinel host” for the detection of *X. fastidiosa* ST53 at low prevalences of host infection.

Although the short latent period and the reliable localisation of the pathogen in infected vectors suggests that vector surveillance could reduce the long detection lags and low diagnostic sensitivities which constrain host surveillance, very little data were available on how the prevalence in vectors relates to that in hosts during early stage spread. We therefore estimate this using a mechanistic model, created to reflect the dynamics of the main vector of *X. fastidiosa, P. spumarius*, in Apulia. Our model predicts that the apparent prevalence in these vectors would be around four times higher than that in hosts in the early stages of infection, which would mean that lower sampling rates would be required in vectors than in hosts during early pathogen spread to reliably sample infected individuals. This high vector prevalence is supported by the observed rapid spread of *X. fastidiosa* ST53 between Apulian olive trees by *P. spumarius* despite the limited transmission window each year (when adults are present and feeding on olive). However, further data on both host and vector infection during early stage spread for future pathogen incursions are urgently needed to verify these conclusions.

Although we assume that the diagnostic sensitivity of vector PCR is reasonably high (resulting from recent advances in vector PCR testing diagnostic methods and protocols (Cruaud et al., 2018; Cunty A., 2019)), PCR tests are expensive and laborious to undertake. As a result, we found that although fewer vectors need to be sampled than hosts, the costs of testing these individually was higher than that of hosts. However, assuming that the performance of PCR is unaffected when five insects are pooled together (as has been suggested (EPPO, 2019)), the total costs of vector testing are lower than those of visual inspection (Figure 6B). Given that the costs and performance of these molecular tests are likely to improve over time, these results suggest that vector testing also offers great future potential for improving the early detection of *X. fastidiosa*.

Further work will be needed to develop appropriate responses to detection in vectors, given that it is not possible to perform repeat confirmatory tests (as is possible with host trees) and that less information can be gained on the spatial distribution of infection. As a result, vector surveys alone are currently not considered sufficient to determine the *X. fastidiosa* status of any area in the European Union (EFSA et al., 2020a). Whilst our method can identify the value of vector surveillance from a scientific perspective, the most appropriate response to a positive detection is best considered by decision makers.

## Conclusions

The rapid rate of spread of *X. fastidiosa* ST53 in olive and the considerable delay between infection and the development of symptoms makes visual inspection less able to identify low prevalences of infection required for effective detection surveys. Whilst molecular tests can reduce the delay before infection can be detected, the relatively low diagnostic sensitivity and high costs of these tests mean that they are unlikely to outperform visual inspection in the field. However, the combination of a short interval between infection and reliable detection and the high initial prevalences of infection amongst the insect vectors responsible for pathogen spread means that vector sampling offers great potential for a sustainable and effective surveillance strategy. Whilst individual testing of vectors is unlikely to currently be a cost-effective alternative to visual inspection, costs can be substantially reduced when insects are pooled together for testing.

## Supporting information

Supplementary Information A: Estimating the performance of absence sampling

Supplementary Information B: Modelling the spread of Xylella fastidiosa between hosts and vectors

Supplementary Information C: Estimating the host prevalence when sampling hosts and vectors

## Acknowledgements

The present work has been funded by Horizon 2020 Project No. 727987 XF-ACTORS (Xylella Fastidiosa Active Containment Through a Multidisciplinary-Oriented Research Strategy)

## Competing Interests

The authors declare no competing interests.

## Author contributions

Initial concept: S.P.; F.v.d.B.

Concept development: F.v.d.B.; A.J.M.; S.P.; Y.B.

Model creation: A.J.M.; F.v.d.B.; Y.B.

Manuscript writing: A.J.M.; S.P.; F.v.d.B.; Y. B.

## Notes

### Competing Interest Statement

The authors have declared no competing interest.

