## Supplementary Information A: Estimating the performance of absence sampling for "Out of sight: Surveillance strategies for emerging vectored plant pathogens"

#### Work so far

Our previous work (Bourhis et al., 2019) has demonstrated that the $x^{th}$ percentile of the prevalence distribution when $N$ hosts are inspected and none are found to be infected is given as:

$$q_A = \left( \frac{-\ln\left(1 - \left(\frac{x}{100}\right)\right)}{(N+1)} \right) \approx \left( \frac{-\ln\left(1 - \left(\frac{x}{100}\right)\right)}{N} \right)$$

We consider here the apparent (i.e. detectable) prevalence $(q_A)$ rather than the true prevalence $(q_T)$. We show later how the latter can be estimated from this.

We have also shown that this can be expanded to any number $(K)$ of equally spaced monitoring rounds, with $N$ hosts inspected at a time interval of $\Delta$:

$$q_A = \left( \frac{-\ln\left(1 - \left(\frac{x}{100}\right)\right)}{A} \right)$$

Where $A$ is estimated as:

$$A = N\left(\frac{\lambda - \lambda^{1-K}}{(\lambda - 1)}\right)$$

$$\lambda = \exp(r\Delta)$$

#### Incorporating imperfect test performance

We consider here how to include an imperfect test in this framework. Although we include both imperfect diagnostic sensitivity and diagnostic specificity here, in most absence sampling situations it would be reasonable to assume a perfect specificity (as any suspected positive cases would likely undergo a confirmatory test).

We derive the probability of a false negative $(\theta_n)$ from the diagnostic sensitivity:

$$\theta_{fn} = 1 - Se$$

And the probability of a false positive $\left(\theta_{fp}\right)$ from the diagnostic specificity:

$$\theta_{fp} = 1 - Sp$$

Representing the apparent prevalence of infection with $q$, we can estimate the probability of different outcomes:

| | | Reality | |
|---|---|---|---|
| | | $1-q$ | $q$ |
| | | Undetectable | Detectable |
| Observation | Negative | $(1-q)(1-\theta_p)$ | $q\theta_n$ |
| | Positive | $(1-q)\theta_p$ | $q(1-\theta_n)$ |

From this, the probability to observe a detectable infected individual is $p_1 = (1-q)\theta_{fp} + q(1-\theta_{fn})$, and the probability to observe an uninfected or undetectable host is $p_0 = (1-q)\left(1-\theta_{fp}\right) + q\theta_{fn}$

The probability that $N$ hosts are observed to be uninfected at any given sampling point is:

$$\left((1-q)\left(1-\theta_{fp}\right)+q\theta_{fn}\right)^N = \left(1-\theta_{fp}\right)^N \left(1-q+\frac{\theta_n}{1-\theta_{fp}}q\right)^N =$$

$$= \left(1-\theta_p\right)^N \left(1-\left(1-\frac{\theta_{fn}}{1-\theta_{fp}}\right)q\right)^N$$

This corresponds to the probability that a host is observed to be uninfected, raised to the power of the sample size. To account for repeated sampling rounds, we need to consider how the apparent prevalence, $q$, changes over time (since this affects the second term in the equation above). If we assume that the apparent prevalence is sufficiently low for exponential growth to be a reasonable approximation, we can introduce $\lambda = \exp(r\varDelta)$, where $r$ is the epidemic growth rate, $\varDelta$ is the sampling interval, and $\lambda$ is therefore the relative increase in the apparent prevalence over this time period (Bourhis et al., 2019). This means that for any prevalence at a given time ($q_t$), the prevalence at the previous sampling point $q_{(t-\Delta)}$ can be estimated as $\frac{q_t}{\lambda}$, or $q_t\lambda^{-1}$. We can keep on repeating this process in order to estimate a general scaling factor for any given number of sampling rounds prior to the current one.

Consider a monitoring program where $K$ monitoring rounds have been done (although in reality, $K$ will generally be unknown). To make the method as generally applicable as possible, we allow both the number of samples and the time interval between subsequent sampling rounds to vary. Starting by denoting the current sampling around as ($i = 1$), we count previous sampling rounds as increasing values of $i$. We assume that in monitoring round $i$, $N_i$ samples were taken, and that the time interval between any monitoring round ($i$) and the subsequent one ($i-1$) is denoted as $\varDelta_i$. We can estimate the total product of prevalence scaling factors up until sampling round $i$ as follows:

| | | | | | | | | | | |
|---|---|---|---|---|---|---|---|---|---|---|
| | First round | | | | | | | | | Latest round |

| Number of monitoring rounds | $K$ | $K-1$ | $K-2$ | … | $i$ | … | 4 | 3 | 2 | 1 |
|---|---|---|---|---|---|---|---|---|---|---|
| $Z_i$ | | | | | * | | $\lambda_3^{-1}\lambda_2^{-1}\lambda_1^{-1}\lambda_0^{-1}$ | $\lambda_2^{-1}\lambda_1^{-1}\lambda_0^{-1}$ | $\lambda_1^{-1}\lambda_0^{-1}$ | $\lambda_0^{-1} = 1$ |
| $q_i$ | | | | | | | $q\lambda_3^{-1}\lambda_2^{-1}\lambda_1^{-1}\lambda_0^{-1}$ | $q\lambda_2^{-1}\lambda_1^{-1}\lambda_0^{-1}$ | $q\lambda_1^{-1}\lambda_0^{-1}$ | $q\lambda_0^{-1}$ |

$$\lambda_{i-1}^{-1}\lambda_{i-2}^{-1}\lambda_{i-3}^{-1}\ldots\ldots\lambda_2^{-1}\lambda_1^{-1}\lambda_0^{-1} = \prod_{j=1}^{i}\lambda_{(j-1)}^{-1}$$

We can now convert this back to the original formulation, giving the following:

$$e^{-r\sum_{j=1}^{i}\Delta_{j-1}} = Z_i$$

Where $\Delta_0 = 0$, and therefore $\lambda_0^{-1} = e^0 = 1$ when $i = 1$ (i.e. at the most recent sampling round).

Here, $Z_i$ is the multiplication factor with which the apparent prevalence in monitoring round $i$ is smaller than in the last monitoring round, $i+1$. The probability that no infections are observed in all $K$ monitoring rounds when the apparent prevalence at the most recent sampling round is $q$ is therefore calculated as:

$$P(no\ positives|q) = \prod_{i=1}^{K}\left(1-\theta_{fp}\right)^{N_i}\left(1-\left(1-\frac{\theta_{fn}}{1-\theta_{fp}}\right)Z_i q\right)^{N_i}$$

We can estimate the apparent prevalence given that no positive detections were made using Bayes law:

$$P(q|no\ positives) = \frac{\tilde{P}(q)\ \prod_{i=1}^{K}\left(1-\theta_{fp}\right)^{N_i}\left(1-\left(1-\frac{\theta_{fn}}{1-\theta_{fp}}\right)Z_i q\right)^{N_i}}{\int_0^1 \tilde{P}(q)\ \prod_{i=1}^{K}\left(1-\theta_{fp}\right)^{N_i}\left(1-\left(1-\frac{\theta_{fn}}{1-\theta_{fp}}\right)Z_i q\right)^{N_i} dq}$$

Assuming we have no prior information, we assume $\tilde{P}(q) = Uniform(0,1)$. Further approximating gives us:

$$\left(1-\theta_{fp}\right)^{N_i}\left(1-\left(1-\frac{\theta_{fn}}{1-\theta_{fp}}\right)Z_i q\right)^{N_i} \approx \left(1-\theta_{fp}\right)^{N_i}\left(e^{-\left(1-\frac{\theta_{fn}}{1-\theta_{fp}}\right)Z_i q}\right)^{N_i}$$

Using the power rule $(e^a)^b = e^{ab}$, we can simplify this further:

$$\left(1-\theta_{fp}\right)^{N_i}\left(e^{-\left(1-\frac{\theta_{fn}}{1-\theta_{fp}}\right)Z_i q}\right)^{N_i} = \left(1-\theta_{fp}\right)^{N_i}e^{-\left(1-\frac{\theta_{fn}}{1-\theta_{fp}}\right)Z_i N_i q}$$

Therefore, we get:

$$\prod_{i=1}^{K}\left(1-\theta_{fp}\right)^{N_i}\left(1-\left(1-\frac{\theta_{fn}}{1-\theta_{fp}}\right)Z_i q\right)^{N_i} = \prod_{i=1}^{K}\left(1-\theta_{fp}\right)^{N_i} e^{-\left(1-\frac{\theta_{fn}}{1-\theta_{fp}}\right)Z_i N_i q} =$$

$$= \prod_{i=1}^{K}\left(1-\theta_{fp}\right)^{N_i} e^{-\left(1-\frac{\theta_{fn}}{1-\theta_{fp}}\right)\sum_{i=1}^{K} Z_i N_i q}$$

and thus:

$$P(q|no\ positives) = \frac{\prod_{i=1}^{K}\left(1-\theta_{fp}\right)^{N_i} e^{-\left(1-\frac{\theta_{fn}}{1-\theta_{fp}}\right)\sum_{i=1}^{K} Z_i N_i q}}{\int_0^1 \prod_{i=1}^{K}\left(1-\theta_{fp}\right)^{N_i} e^{-\left(1-\frac{\theta_{fn}}{1-\theta_{fp}}\right)\sum_{i=1}^{K} Z_i N_i q}\, dq} =$$

$$= \frac{e^{-\left(1-\frac{\theta_{fn}}{1-\theta_{fp}}\right)\sum_{i=1}^{K} Z_i N_i q}}{\int_0^1 e^{-\left(1-\frac{\theta_{fn}}{1-\theta_{fp}}\right)\sum_{i=1}^{K} Z_i N_i q}\, dq}$$

As we are interested in very small values of $q$ only, the density for $q$ a bit larger than very small is virtually zero. This implies that integrating from 0 to 1 is approximately the same as integrating from 0 to infinity.

We therefore finally get:

$$P(q|no\ positives) = \left(1-\frac{\theta_{fn}}{1-\theta_{fp}}\right)\sum_{i=1}^{K} Z_i N_i\ e^{-\left(1-\frac{\theta_{fn}}{1-\theta_{fp}}\right)\sum_{i=1}^{K} Z_i N_i q}$$

Which can also be represented as:

$$P(q|no\ positives) = A\, e^{-Aq}$$

$$A = \left(1-\frac{\theta_{fn}}{1-\theta_{fp}}\right)\sum_{i=1}^{K} Z_i N_i$$

In line with our previous work, this is an exponential distribution.

For constant sample number, $N_i = N$, and sample interval, $\Delta_i = \Delta$, we get

$$A = \left(1-\frac{\theta_{fn}}{1-\theta_{fp}}\right) N \sum_{i=1}^{K}\left(\lambda^{-1}\right)^{i-1}$$

$$\sum_{i=1}^{K}\left(\lambda^{-1}\right)^{i-1} = \left(\lambda^{-1}\right)^{0} + \left(\lambda^{-1}\right)^{1} + \left(\lambda^{-1}\right)^{2} + \cdots \ldots \ldots \ldots + \left(\lambda^{-1}\right)^{K-1}$$

This can also be considered as the following:

$$S = 1 + \left(\frac{1}{\lambda}\right) + \left(\frac{1}{\lambda^2}\right) + \cdots \ldots \ldots \ldots + \left(\frac{1}{\lambda^{(K-1)}}\right)$$

This is a geometric series, and can be solved as follows:

$$S\left(\frac{1}{\lambda}\right) = \left(\frac{1}{\lambda}\right) + \left(\frac{1}{\lambda^2}\right) + \left(\frac{1}{\lambda^3}\right) \dots\dots\dots\dots + \left(\frac{1}{\lambda^K}\right)$$

Subtracting this from the first formulation gives:

$$S - S\left(\frac{1}{\lambda}\right) = 1 - \left(\frac{1}{\lambda^K}\right)$$

$$= S\left(1 - \lambda^{-1}\right) = 1 - \lambda^{-K}$$

Which gives:

$$S = \frac{1 - \lambda^{-K}}{1 - \lambda^{-1}} = \frac{\lambda\left(1 - \lambda^{-K}\right)}{\lambda - 1}$$

Which can also be presented as:

$$S = \frac{1 - \left(\lambda^{-1}\right)^K}{1 - \left(\lambda^{-1}\right)} = \frac{\lambda - \lambda^{1-K}}{\lambda - 1}$$

So

$$A = \left(1 - \frac{\theta_{fn}}{1 - \theta_{fp}}\right) N \, \frac{\lambda - \lambda^{1-K}}{\lambda - 1}$$

However, we still don't know what $K$ is (and are unlikely to know).

As $K$ increases, the denominator in the series gets increasingly large (meaning that $1 - \lambda^{-K} \approx 1$), allowing us to treat this as an infinite series. Using the same approach as above, but with an infinite series each time gives us the following:

$$S - S\left(\frac{1}{\lambda}\right) = 1$$

$$S\left(1 - \lambda^{-1}\right) = 1$$

$$S = \frac{1}{1 - \lambda^{-1}} = \frac{\lambda}{\lambda - 1}$$

This therefore gives us the following final formulation (given that $\Delta$ and $N$ are fixed and that $K$ is reasonably large – i.e. that the sampling program has been in place for a long time, which I imagine is a reasonable expectation):

$$P(q|no\ positives) = \ Ae^{-Aq}$$

$$A = \left(1 - \frac{\theta_{fn}}{1 - \theta_{fp}}\right)\left(\frac{\lambda}{\lambda - 1}\right) N$$

The $x^{th}$ percentile of this distribution $(q_x)$ is estimated for the first round of sampling as:

$$q_x = \frac{-\ln\left(1 - {}^{x}/_{100}\right)}{\left(1 - \frac{\theta_{fn}}{1-\theta_{fp}}\right) N}$$

And for all subsequent rounds of sampling as:

$$q_x = \left(\frac{-\ln\left(1 - {}^{x}/_{100}\right)}{\left(1 - \frac{\theta_{fn}}{1-\theta_{fp}}\right) N}\right)\left(1 - \frac{1}{\lambda}\right) = \left(\frac{-\ln\left(1 - {}^{x}/_{100}\right)}{\left(1 - \frac{\theta_{fn}}{1-\theta_{fp}}\right) N}\right)(1 - \mathrm{e}^{-r\Delta})$$

Under the assumption of perfect diagnostic specificity, $\theta_{fp} = 0$ and we can simplify this down further. Estimating the minimum required sample size to detect at a detectable prevalence of $q_x$ and using $Se$ to indicate the diagnostic sensitivity:

1st monitoring round:

$$N > \left(\frac{-\ln\left(1 - {}^{x}/_{100}\right)}{Se}\right)\left(\frac{1}{q_x}\right)$$

All other monitoring rounds:

$$N > \left(\frac{-\ln\left(1 - {}^{x}/_{100}\right)}{Se}\right)\left(\frac{1}{q_x}\right)\ (1 - e^{-r\Delta})$$

### Accounting for the detection lag

We have shown that the $x^{th}$ percentile of the prevalence distribution after a single round of sampling with no detections is estimated as:

$$q_A = \left(\frac{-\ln\left(1 - {}^{x}/_{100}\right)}{SeN}\right)$$

If we assume that the true prevalence is still low enough for its growth to be increasing exponentially, its relationship to the apparent prevalence is as follows for a detection lag of $\delta$ and a growth rate of $r$:

$$q_T = q_A \cdot e^{r\delta}$$

The true prevalence is then estimated as:

$$q_T = \left(\frac{-\ln\left(1 - {}^{x}/_{100}\right)}{SeN}\right)e^{r\delta}$$

And therefore:

$$N > \left(\frac{-\ln\left(1 - {}^{x}/_{100}\right)}{Se}\right)\left(\frac{e^{r\delta}}{q_T}\right)$$

However, when δ and $r$ are high, the true prevalence may be increasing logistically rather than exponentially:

$$q_T = \left( \frac{q_A \cdot e^{r\delta}}{(1 + q_A(e^{r\delta} - 1))} \right)$$

Combining these gives:

$$q_T = \left( \frac{-\ln\left(1 - {}^{x}/_{100}\right) e^{r\delta}}{\ln\left(1 - {}^{x}/_{100}\right)(1 - e^{r\delta}) + SeN} \right)$$

And therefore:

$$N > \left( \frac{-\ln\left(1 - {}^{x}/_{100}\right)}{Se} \right) \left( 1 + e^{r\delta} \left( \frac{q_T - 1}{q_T} \right) \right)$$

For multiple sampling rounds, the $x^{th}$ percentile of the prevalence distribution after a single round of sampling with no detections is estimated as:

$$q_A = \left( \frac{-\ln\left(1 - {}^{x}/_{100}\right)}{SeN} \right) (1 - \mathrm{e}^{-r\Delta})$$

$$q_T = \left( \frac{-\ln\left(1 - {}^{x}/_{100}\right) e^{r\delta}(\mathrm{e}^{r\Delta} - 1)}{\left(-\ln\left(1 - {}^{x}/_{100}\right)(e^{r\delta} - 1)(e^{r\Delta} - 1) + SeNe^{r\Delta}\right)} \right)$$

And therefore:

$$N = \left( \frac{-\ln\left(1 - {}^{x}/_{100}\right)}{Se} \right) \left( 1 + e^{r\delta} \left( \frac{q_T - 1}{q_T} \right) \right) (1 - e^{-r\Delta})$$

### Comparing different detection methods

We assess the relative value of two detection methods in order to detect at the same threshold incidence (Mastin et al., 2019). Using again the example of the first monitoring round only:

For visual assessment:

$$q_T = \left( \frac{-\ln\left(1 - {}^{x}/_{100}\right) e^{r\delta_1}}{\ln\left(1 - {}^{x}/_{100}\right)(1 - e^{r\delta_1}) + Se_1 N_1} \right)$$

For an alternative detection method:

$$q_T = \left( \frac{-\ln\left(1 - {x}/{100}\right) e^{r\delta_2}}{\ln\left(1 - {x}/{100}\right)(1 - e^{r\delta_2}) + Se_2 N_2} \right)$$

In order to detect at the same true prevalence, the two detection methods are equivalent when the following is true:

$$\frac{Se_1 N_1 + \ln\left(1 - {x}/{100}\right)}{e^{r\delta_1}} = \frac{Se_2 N_2 + \ln\left(1 - {x}/{100}\right)}{e^{r\delta_2}}$$

We can consider the minimum total cost of sampling $(C)$ as the product of the sample size and the per-host cost of sampling $(c)$:

$$C = Nc$$

$$C_1 = \left( \frac{-\ln\left(1 - {x}/{100}\right)}{Se_1} \right)\left(1 + e^{r\delta_1}\left(\frac{q_T - 1}{q_T}\right)\right) c_1$$

$$C_2 = \left( \frac{-\ln\left(1 - {x}/{100}\right)}{Se_2} \right)\left(1 + e^{r\delta_2}\left(\frac{q_T - 1}{q_T}\right)\right) c_2$$

From this, we can estimate the relative total cost of visual detection as follows:

$$\frac{C_1}{C_2} = \frac{\left(\frac{c_1}{Se_1}\right)\left(1 + e^{r\delta_1}\left(\frac{q_T - 1}{q_T}\right)\right)}{\left(\frac{c_2}{Se_2}\right)\left(1 + e^{r\delta_2}\left(\frac{q_T - 1}{q_T}\right)\right)}$$

This demonstrates that the relative total cost is not affected by the confidence level, although it is affected by the true prevalence threshold under the assumption of logistic growth (our previous work has shown that this is not the case under the assumption of exponential growth).

Alternatively, the maximum relative per-host cost of visual inspection for a fixed total cost can be estimated as:

$$\left(\frac{c_1}{c_2}\right) < \frac{\left(\frac{\left(1 + e^{r\delta_2}\left(\frac{q_T - 1}{q_T}\right)\right)}{Se_2}\right)}{\left(\frac{\left(1 + e^{r\delta_1}\left(\frac{q_T - 1}{q_T}\right)\right)}{Se_1}\right)}$$

For multiple sampling, the total cost is simply:

$$C_1 = \left( \frac{-\ln\left(1 - {x}/{100}\right)}{Se_1} \right)\left(1 + e^{r\delta_1}\left(\frac{q_T - 1}{q_T}\right)\right)(1 - e^{-r\Delta}) c_1$$

$$C_2 = \left(\frac{-\ln\left(1 - {}^{x}\!/_{100}\right)}{Se_2}\right)\left(1 + e^{r\delta_2}\left(\frac{q_T - 1}{q_T}\right)\right)(1 - e^{-r\Delta})c_2$$

And the cost ratio is equal to that for a single sample, demonstrating that the relative value of the diagnostic is not affected by the number of samples.

$$\frac{C_1}{C_2} = \frac{\left(\frac{c_1}{Se_1}\right)\left(1 + e^{r\delta_1}\left(\frac{q_T - 1}{q_T}\right)\right)}{\left(\frac{c_2}{Se_2}\right)\left(1 + e^{r\delta_2}\left(\frac{q_T - 1}{q_T}\right)\right)}$$
