## Supplementary Information B: Modelling the spread of Xylella fastidiosa between hosts and vectors for "Out of sight: Surveillance strategies for emerging vectored plant pathogens"

We describe here the formulation of our model of host and vector transmission. Importantly, this model explicitly incorporates seasonality and we therefore consider the state of the system for different Julian days $(t)$ of the year, as well as between years. We denote the density of uninfected and infected vectors on day $t$ of a given year as $X(t)$ and $Y(t)$, respectively. As well as uninfected $\left(H(t)\right)$ and infectious $(I)$ hosts, we capture the host latent period by also considering those hosts which are infected but not yet infectious $\left(E(t)\right)$. Whilst the density of trees in the $E(t)$ compartment varies over each growing season (in relation to the density of infected vectors), we assume that the density of trees in the $I$ compartment only varies between growing seasons. However, since uninfected vectors become infected by feeding on these infectious olive trees and these drive the transition from $H(t)$ to $E(t)$, hosts in the $I$ compartment do not effectively become infectious until the first day of adult vector emergence (when transmission to other hosts can start to take place).

#### Vector dynamics

We start by considering how the total density of adult vectors $(P(t))$ changes over time. Since we assume that vectors are not affected by infection with *X. fastidiosa*, these population dynamics can be described independently of infection status. We can assume that the rate of increase in vector density is the product of the net population growth rate of the adult population at time $t$ $\left(g(t)\right)$ and the density of adults:

$$\frac{dP(t)}{dt} = g(t)P(t)$$

We assume that adults emerge at rate $g_0$ and that the emergence rate decreases with time. We also assume that the adult death rate increases with time due to adult age and the increase in adverse weather conditions later in the year. We can include both the decrease in emergence rate and the increase in the death rate in a single parameter, $a$:

$$g(t) = (g_0 - at)$$

Including this in the differential equation above and solving gives:

$$P(t) = P_0 \, e^{\left(g_0 t - \frac{a}{2} t^2\right)}$$

Where $P_0$ reflects the initial density of adults emerging. Since adult vectors do not emerge at the start of the year, we need to consider $t$ in relation to the day at which they start to emerge $(t_{em})$:

$$P(t) = P_0 \, e^{g_0 (t - t_{em}) - \frac{a}{2}(t - t_{em})^2}$$

We now consider the density of infected vectors. This would be expected to change according to the density of uninfected vectors, the density of infected hosts, and the density of infected vectors as follows:

$$\frac{dY(t)}{dt} = \beta X(t)I - \mu(t)Y(t)$$

Where $\beta$ is the rate at which vectors acquire the bacterium and $\mu(t)$ is the vector death rate on day $t$. As the total adult vector population, $P(t)$, is the sum of the uninfected and infected sub-populations on day $t$, we can present this equation in terms of $Y(t)$:

$$\frac{dY(t)}{dt} = \beta\big(P(t) - Y(t)\big)I - \mu(t)Y(t)$$

The proportion of the vector population which is infected (the vector prevalence, $q_V(t)$) can then be estimated as the ratio of the infected vector density and total vector density:

$$q_V(t) = \frac{Y(t)}{P(t)}$$

Taking derivatives:

$$\frac{dq_V(t)}{dt} = \frac{P(t)\frac{dY(t)}{dt} - \frac{dP(t)}{dt}Y(t)}{P^2(t)}$$

Substituting the various terms gives:

$$\frac{dq_V(t)}{dt} = \frac{\beta\big(P(t) - Y(t)\big)I - \mu(t)Y(t)}{P(t)} - q_V(t)\frac{dP(t)/dt}{P(t)}$$

$$\frac{dq_V(t)}{dt} = \beta I - \beta I q_V(t) - \mu(t)q_V(t) - q_V(t)(g_0 - at)$$

$$\frac{dq_V(t)}{dt} = \beta I - \big(\beta I + \mu(t) + (g_0 - at)\big)q_V(t)$$

Assuming $at = \mu(t)$:

$$\frac{dq_V(t)}{dt} = \beta I - (\beta I + g_0)q_V(t)$$

This has solution:

$$q_V(t) = \frac{\beta I}{\beta I + g_0}\left(1 - e^{-(\beta I + g_0)t}\right)$$

We now need to consider that vectors cannot start to acquire the bacteria until after adult emergence. Denoting the day at which acquisition starts as $t_{in}$, we have:

$$q_V(t) = \frac{\beta I}{\beta I + g_0}\left(1 - e^{-(\beta I + g_0)(t - t_{in})}\right)$$

The density of infected vectors on Julian day $t$, assuming emergence from day $t_{em}$and bacterial acquisition from day $t_{in}$, is thus given by:

$$Y(t) = q_V(t)P(t) = P_0\left(\frac{\beta I}{\beta I + g_0}\right)\left(1 - e^{-(\beta I + g_0)(t - t_{in})}\right)e^{g_0(t - t_{em}) - \frac{a}{2}(t - t_{em})^2}$$

### Host dynamics

We now consider the dynamics of host infection. We assume that the total density of host trees $(D)$ does not change over our timescale of interest, and comprises uninfected $\big(H(t)\big)$ and infected exposed $\big(E(t)\big)$ hosts, which change within a given year, and infected infectious hosts $(I)$ which only change between years:

$$D = H(t) + E(t) + I$$

Over the course of the year, the density of newly infected (not yet infectious or detectable) hosts changes as:

$$\frac{dE(t)}{dt} = \alpha H(t) Y(t)$$

Where $\alpha$ is the rate of host inoculation by vectors. Since $H(t) = D - E(t) - I$:

$$\frac{dE(t)}{dt} = \alpha(D - E(t) - I)Y(t)$$

Noting that infected vectors are only present from day $t_{in}$ onwards, this is solved as:

$$-\ln(D - E(t) - I) + C = \alpha \int_{t_{in}}^{t} Y(x)dx \text{ for } t \geq t_{in}$$

The integration constant $C$ is found by substituting $t = 0$:

$$C = \ln(D - I)$$

Reformulating to estimate $E(t)$:

$$\ln(D - E(t) - I) = \ln(D - I) - \alpha \int_{t_{in}}^{t} Y(x)dx$$

$$(D - E(t) - I) = (D - I)\exp\left(-\alpha \int_{t_{in}}^{t} Y(x)dx\right)$$

$$E(t) = (D - I)\left(1 - \exp\left(-\alpha \int_{t_{in}}^{t} Y(x)dx\right)\right)$$

To calculate the total number of new infections by the end of the year, we need to evaluate the integral of the infected vector density $Y(t)$ between $t = t_{in}$ and $t = 365$:

$$\int_{t_{in}}^{365} Y(x)dx = P_0\left(\frac{\beta I}{\beta I + g_0}\right) \int_{t_{in}}^{365} \left(1 - e^{-(\beta I + g_0)(t - t_{in})}\right) e^{g_0(t - t_{im}) - \frac{a}{2}(t - t_{im})^2}\, dt$$

This integral can be interpreted as the total density of infected-vector-days over the year, and therefore $\exp\left(-\alpha \int_{t_{in}}^{t} Y(x)dx\right)$ is the probability that a host does not get infected during that year (meaning that $\left(1 - \exp\left(-\alpha \int_{t_{in}}^{t} Y(x)dx\right)\right)$ is the probability of at least one infection over the year). The total density of hosts which become newly infectious $(I)$ each year is assumed to be equal to the total density of hosts which become newly infected over the course of the previous year. Presenting this as a difference equation:

$$I_{i+1} = I_i + (D - I_i)\left(1 - \exp\left(-\alpha \int_{t_{in}}^{t=365} Y(x)dx\right)\right)$$

Although $I$ does not change over the course of the year, spread from these infectious hosts does not commence until vectors emerge and start to acquire the bacteria (on day $t_{in}$). As a result, this approach assumes a latent period of between $t_{in}$ and 365 days, which is in line with current knowledge. It also assumes that the latent and presymptomatic periods are equal (meaning that symptoms would start to develop at the same time as a host becomes infectious), and therefore $I_i$ describes the density of symptomatic hosts as well as the density of infectious hosts.

### Parameter estimation

We start by considering the rate of adult vector emergence ($g_0$). Aggregating by month and taking the average vector prevalence over time from three studies:

| month | Julian day | Ben Mossa | Cornara JEE | Cornara JAE | Mean of 3 |
|---|---|---|---|---|---|
| 1 | 30 | (0.25) | -- | | |
| 2 | 60 | -- | -- | | |
| 3 | 90 | 0 | -- | | |
| 4 | 120 | 0 | -- | | |
| 5 | 150 | 0 | 0.27 | 0.178 | 0.178 |
| 6 | 180 | 0.31 | 0.55 | 0.476 | 0.476 |
| 7 | 210 | 0.19 | 0.51 | 0.414 | 0.414 |
| 8 | 240 | 0.20 | 1.0 | 0.569 | 0.569 |
| 9 | 270 | 0.14 | 0.80 | 0.492 | 0.492 |
| 10 | 300 | 0.31 | 0.65 | 0.4579 | 0.458 |
| 11 | 330 | 0.1 | -- | | |
| 12 | 360 | 0 | -- | | |

Fitting these data to the following equation with $t_{in} = 150$ (which represents the time at which infected vectors were first observed):

$$q_V(t) = \frac{\beta I}{\beta I + g_0}\left(1 - e^{-(\beta I + g_0)(t - t_{in})}\right)$$

We get:

$$\frac{\beta I}{\beta I + g_0} = 0.484 \pm 0.05339$$
$$(\beta I + g_0) = 0.1258 \pm 0.3557$$
$$R^2 = 0.876$$

Therefore:

$$\beta I = \frac{0.484}{0.1258} = 0.060887$$

$$g_0 = 0.1258 - 0.060887 = 0.064913$$

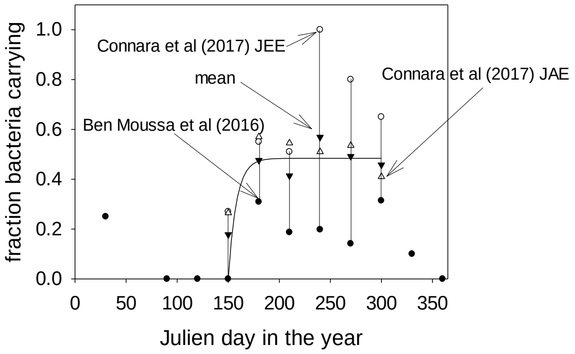

We next consider the initial density of adults emerging $(P_0)$ and the rate of decline in the numbers of adults with time $(a)$. Unfortunately, we do not have any estimates of the absolute density of vectors over time, only estimates of the relative changes in density. We therefore aggregated the density data from two studies by month and rescaled these estimates to lie within the range [0,1]:

| Julian day | Ben Moussa et al (2016) | Cornara et al (2017) | Mean |
|---|---|---|---|
| 30 | 0.026936 | -- | |
| 60 | 0 | -- | |
| 90 | 3.37E-03 | -- | |
| 120 | 0.16835 | -- | |
| 150 | 0.215488 | 0.513514 | 0.365 |
| 180 | 0.37037 | 0.486486 | 0.428 |
| 210 | 0.521886 | 1 | 0.761 |
| 240 | 0.680135 | 0.324324 | 0.502 |
| 270 | 1 | 0.27027 | 0.635 |
| 300 | 0.171717 | 0.040541 | 0.106 |
| 330 | 0.03367 | 0 | 0.016 |
| 360 | 0.023569 | 0 | 0.011 |

We can fit these data to the following equation with $t_{em} = 80$ (which represents the time at which adult vectors were first observed in (Ben Moussa et al., 2016)) and the $g_0$ estimate derived above:

$$P(t) = C_1\, e^{g_0(t-t_{em}) - \frac{a}{2}(t-t_{em})^2}$$

Note that we use $C_1$ rather than $P_0$ as we are using relative density data and the interpretation is different.

We get:

$$C_1 = 0.007265 \pm 0.001922$$

$$\frac{a}{2} = 0.0002287 \pm 0.00001165$$

$$R^2 = 0.812$$

Therefore:

$$a = 0.0004574$$

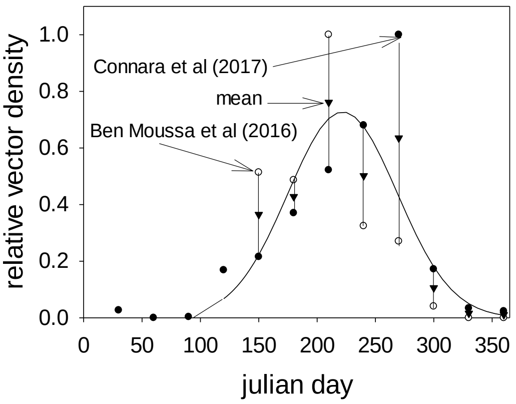

Note that this is relative densities. So we need to rescale $C_1$ to get a value that represents the actual population density. However, this parameter currently reflects both the initial population density of adults emerging into the olive trees $(P_0)$ and a scaling factor for $g_0$ and $a$. To illustrate this, for the case where the parameter now represents the actual densities, we write $C = P_0 \ln(Z)$. We then get

$$P(t) = P_0\, e(Z)\, e^{g_0(t-t_{em})-\frac{a}{2}(t-t_{em})^2} = P_0\, e^{Z\left(g_0(t-t_{em})-\frac{a}{2}(t-t_{em})^2\right)}$$

We need to introduce a valid $P_0$ to get real densities, which should be such that the peak value of $P(t)$ is the value observed in the field. The peak value of the fitted curve is at day 225 and is 0.725, and the expected peak adult density in the field (from expert opinion) is 20 adults /m$^2$. Therefore, we multiply $C_1$ by $P_0 = \frac{20}{0.725} = 27.59$.

We now consider the rate of pathogen acquisition by vectors $(\beta)$. We find that the fraction of infected hosts $\left(\frac{I}{D}\right)$ in the areas covered by the vector studies (Ben Moussa et al., 2016; Cornara, Cavalieri, et al., 2017; Cornara, Saponari, et al., 2017) is 0.23. Therefore:

$$\frac{\beta I}{\left(\frac{I}{D}\right)} = \frac{0.060887}{0.23} = 0.26472608 = \beta D$$

$$\beta = \frac{\beta D}{D} = \frac{0.26472608}{0.01234568} = 21.4428$$

Finally, we estimate the rate of inoculation of hosts by vectors $(\alpha)$.

When $I_i$ is very small, then $A$ is very small, and we can linearise our host difference equation as follows:

$$I_{i+1} = I_i + (D - I_i)\left(1 - \exp\left(-\alpha A P_0\left(\frac{\beta I_i}{\beta I_i + g_0}\right)\right)\right) \approx I_i + D\alpha \left|\frac{dA}{dI_i}\right|_{I_i=0} I_i$$

Where:

$$\left|\frac{dA}{dI_i}\right|_{I_i=0} = P_0\left(\frac{\beta}{g_0}\right)\int_{t_{in}}^{365}(1-e^{-g_0 t})e^{g_0 t-\frac{a}{2}t^2}dt = P_0\left(\frac{\beta}{g_0}\right)A$$

The integral is the value of $A$ at $I = 0$. Then:

$$\left|\frac{d\ A}{d\ I_i}\right|_{I_i=0} = P_0\left(\frac{\beta}{g_0}\right)\ 13241.238 = \ 120{,}662{,}122.912$$

The largest eigenvalue is $\lambda\ = 1 + D\alpha\ \left|\frac{d\ A}{d\ I_i}\right|_{I_i=0}$.

This can also be estimated from the exponential growth rate, $r = 0.0122$. Using the estimated $r$, $\lambda\ =\ e^{r\times 365} =\ e^{0.0122\times 365} = 85.9$. However, this is an overestimate as infected vectors are only present for $365 - t_{in} = 215$ days in the year. Therefore $\lambda = e^{0.0122\times 215} = 13.78$.

Therefore:

$$\alpha = (\lambda - 1)\frac{1}{D\left|\frac{d\ A}{d\ I_i}\right|_{I_i=0}}$$

$$\frac{1}{D\left|\frac{d\ A}{d\ I_i}\right|_{I_i=0}} = 6.71296486215 \times 10^{-7}$$

$$\alpha = 8.579169 \times 10^{-6}$$

We can also simplify the host model. Consider the integral expression describing the dynamics of infected vectors:

$$A = \int_{t_{in}}^{365}\left(1 -\ e^{-(\beta I+g_0)(t-t_{in})}\right)\ e^{g_0(t-t_{im})-\frac{a}{2}(t-t_{im})^2}\ dt$$

Using our estimates of $\beta$, $g_0$, $t_{in}$, and $t_{im}$, when $I = 0$, $A = 13241.238$. When $I = 0.01234467$ (i.e. complete host infection), $A = 13901.913$. This represents a relatively small difference in $A$. Rather than calculating the integral dynamically, we therefore taken the mean $(A = 13571.5965)$ as a reasonable approximation.

This gives us a model for $I$ which does not require integration:

$$I_{i+i} =\ I_i + (D - I_i)\left(1 - \exp\left(-\ \alpha A P_0\left(\frac{\beta I_i}{\beta I_i +\ g_0}\right)\right)\right)$$

Where $\alpha A =\ 0.11643$.
