## Supplementary Information C: Estimating the host prevalence when sampling hosts and vectors for "Out of sight: Surveillance strategies for emerging vectored plant pathogens"

#### Summary of approach

Here, we consider how to evaluate the performance of a surveillance scheme in which hosts and/or vectors in an epidemiologically linked system are sampled and inspected/tested and none are found to be infected. We explicitly consider the effect of "detection lag" periods before infection can be detected by first considering the prevalence of detectable hosts $\left(q_{H_A}\right)$ and vectors $\left(q_{V_A}\right)$ (although in practice we do not consider a detection lag in vectors – effectively assuming that infected vectors would be potentially detectable immediately after initial infection), before estimating the associated true prevalence of host infection $\left(q_{H_T}\right)$ in the final stage of analysis.

#### Probability of detection for a single round of sampling

We first consider one monitoring round. During this monitoring round, $N_H$ hosts undergo visual inspection (where $N_H$ may equal 0 if no hosts are inspected). The probability that an inspected host is uninfected (or presymptomatically infected) is $\left(1-q_{H_A}\right)$. Assuming a false positive rate (one minus the diagnostic specificity) of $\theta_{pH}$, the probability that a visited host is uninfected/ presymptomatic and is identified as uninfected is $\left(1-\theta_{pH}\right)\left(1-q_{H_A}\right)$. The probability that a visited host is infected and symptomatic but is identified as uninfected is $\theta_{nH}q_{H_A}$, where $\theta_{nH}$ is the probability of a false negative (one minus the diagnostic sensitivity). Combining these we see that the probability that a sampled host is assessed as uninfected, $P_H$, is given by

$$P_H=\left(1-\theta_{pH}\right)\left(1-q_{H_A}\right)+\theta_{nH}q_{H_A}$$

Sampling $N_H$ hosts, the probability not to find any infected hosts equals ${P_H}^{N_H}$.

Similarly, the probability that a sampled vector is assessed as uninfected, $P_V$, is:

$$P_V=\left(1-\theta_{pV}\right)\left(1-q_{V_A}\right)+\theta_{nV}q_{V_A}$$

Taking $N_V$ of such bulk samples, the probability not to find any bacteria in the samples is ${P_V}^{N_V}$.

For one entire monitoring round the probability not to find the pathogen when the incidence of the host and vector are $q_{H_A}$ and $q_{V_A}$, respectively $\left(P\left(no\ detections|\ q_{H_A},q_{V_A}\right)\right)$ is:

$$P\left(no\ detections|q_{H_A},q_{V_A}\right)={P_H}^{N_H}\,{P_V}^{N_V}$$

#### Probability of detection for multiple rounds of sampling

We now consider multiple rounds of sampling, where the interval between two monitoring rounds, $\Delta$, and the number of host and vector samples taken in a monitoring round, $N_H$ and $N_V$, are constant.

We first assume that the detectable host prevalence grows exponentially (as would be the case if the apparent prevalence is low):

$$q_{H_A}(t) = q0_{H_A}e^{rt}$$

Where $q0_{H_A}$ is the detectable host incidence at time $t = 0$. We take this exponential growth into account and calculate the detectable prevalence of infection for all previous sampling events when the detectable prevalence at the current sample is $q_{H_A}$.

For the current host detectable prevalence, $q_{H_A}$, the detectable prevalence at the previous sampling point (sample 2) would be equal to:

$$q_{H2_A} = q_{H_A}Z_2 = q_{H_A}e^{-r\Delta}$$

And at the sampling point before that (sample 3), it would be:

$$q_{H3_A} = q_{H_A}Z_3 = q_{H_A}e^{-r2\Delta}$$

We can expand this up to sample $i$:

$$q_{Hi_A} = q_{H_A}Z_i = q_{H_A}e^{-r(i-1)\Delta}$$

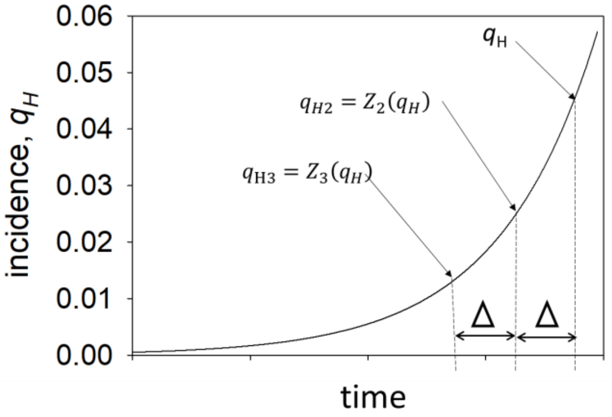


Using this approach, we estimate the detectable prevalences at each previous sampling point from the current detectable prevalence. We can estimate the probability of not obtaining a positive result when $N_H$ samples are taken at any one of these points using the equation presented above:

$$P_{H_i}{}^{N_H} = \left((1-\theta_{pH})(1-q_{H_A}Z_i) + \theta_{nH}q_{H_A}Z_i\right)^{N_H}$$

The probability that in the entire series of $K$ sampling rounds, no infected hosts are detected is then estimated as the product of these for all sampling points:

$$\prod_{i=1}^{K} P_{H_i}{}^{N_H}$$

We can use a similar approach for the vector prevalence. Denoting the equivalent of $Z_i$ for the vector $C_i$, the probability of not detecting any infected vectors in a single sampling round $i$ is given by:

$$P_{V_i}{}^{N_V} = \left((1-\theta_{pV})(1-q_{V_A}C_i)^M + \theta_{nV}\left(1-(1-q_{V_A}C_i)^M\right)\right)^{N_V}$$

The probability that in the entire series of $K$ sampling rounds, no infected vectors are detected is then:

$$\prod_{i=1}^{K} {P_{V_i}}^{N_V}$$

The probability that in the entire monitoring program of $K$ sampling rounds no infected hosts or vectors are found is then:

$$P(no\ detections|q_{H_A}, q_{V_A}) = \prod_{i=1}^{K} {P_{Hi}}^{N_H} {P_{Vi}}^{N_V}$$

To calculate the host and vector incidence when no infected trees or vectors are found we use Bayes theorem relating the required probability density to the probability that the pathogen is not detected given that the frequency of the infection in the target population is $q$, $P(no\ detections|q_{H_A}, q_{V_A})$, by:

$$P(q_{H_A}, q_{V_A}|no\ det) = \frac{P(q_{H_A}, q_{V_A})P(no\ detections|q_{H_A}, q_{V_A})}{\int_0^1 \int_0^1 P(q_{H_A}, q_{V_A})P(no\ detections|q_{H_A}, q_{V_A})dq_{H_A}dq_{V_A}}$$

where the prior distribution of $q_{H_A}$ and $q_{V_A}$, $P(q_{H_A}, q_{V_A})$, is taken to be uninformative as a two dimensional uniform density.

$$P(q_{H_A}, q_{V_A}|no\ det) = \frac{P(no\ detections|q_{H_A}, q_{V_A})}{\int_0^1 \int_0^1 P(no\ detections|q_{H_A}, q_{V_A})\ dq_{H_A}dq_{V_A}}$$

### Incorporating linked host and vector prevalences

The methods described above could be used when sampling from two epidemiologically independent populations, but in the case where the populations are linked (in our case, a pathogen spreading between hosts and vectors), both $Z_i$ and $C_i$ and $q_{H_A}$and $q_{V_A}$ are related to each other. Because $q_{V_A}$ and $q_{H_A}$ are parts of an interconnected dynamic system, it follows that they:

(i) Increase exponentially at the same rate, so $r_H = r_V$. Importantly, this implies that $Z_i = C_i = e^{-r(i-1)\Delta}$

(ii) Exist as a fixed ratio during this initial exponential phase of epidemic growth, so $\frac{q_{V_T}}{q_{H_T}} = \varphi_T$ and $\frac{q_{V_A}}{q_{H_A}} = \varphi_A$.

As we assume that $\varphi_A$ is fixed, the apparent prevalence of infection in hosts $(q_{H_A})$ can be estimated from the apparent prevalence of infection in vectors $(q_{V_A})$, and vice versa. We assume that $q_{H_A}$ and $q_{V_A}$ are both growing exponentially, which is reasonable for the early stages of spread which we are interested in. In order to derive analytic estimates for $\varphi_T$ and $\varphi_A$ in the early phase of the epidemic, we first assume that the number of infected hosts $(I)$ is very low.

We assume that there is no detection lag in vectors, meaning that the apparent and true prevalences are equal:

$$q_V(t) = q_{V_A}(t) = q_{V_T}(t) = \frac{\beta I_i}{\beta I_i + g_0}\left(1 - e^{-(\beta I_i + g_0)(t - t_{in})}\right)$$

The proportion of the host population which is infected (the true host prevalence, $q_{H_T}(t)$) can be estimated as:

$$q_{H_T}(t) = (E(t) + I_i)/D$$

The prevalence of symptomatic hosts (the apparent host prevalence) is:

$$q_{H_A}(i) = I_i/D$$

For small values of *I* (i.e. when the number of infected hosts is low) we get:

$$q_{V_A}(t) \approx \frac{\beta I}{g_0}\left(1 - e^{-g_0(t-t_{in})}\right)$$

Assuming that sampling takes place in the month where the vector is most abundant and the vector prevalence has reached its plateau:

$$q_{V_A} \approx \frac{\beta I}{g_0}$$

The apparent prevalence of infection in hosts is estimated as:

$$q_{H_A} = \frac{I}{D}$$

So the ratio of apparent vector and host prevalences is:

$$\frac{q_{V_A}}{q_{H_A}} = \varphi_A \approx \frac{\frac{\beta I}{g_0}}{\frac{I}{D}} = \frac{D\beta}{g_0}$$

This relationship allows us to reformulate our early equation to relate only to the apparent prevalence in hosts:

$$P_{V_i}{}^{N_V} = \left((1 - \theta_{pV})(1 - \varphi_A q_{H_A} Z_i)^M + \theta_{nV}\left(1 - (1 - \varphi_A q_{H_A} Z_i)^M\right)\right)^{N_V}$$

$$P(no\ detections|q_{H_A}) = \prod_{i=1}^{K}\left((1 - \theta_{pH})(1 - q_{H_A} Z_i) + \theta_{nH} q_{H_A} Z_i\right)^{N_H} \cdot \left((1 - \theta_{pV})(1 - \varphi_A q_{H_A} Z_i)^M + \theta_{nV}\left(1 - (1 - \varphi_A q_{H_A} Z_i)^M\right)\right)^{N_V}$$

And therefore:

$$P(q_{H_A}|no\ det) = \frac{P(no\ detections|q_{H_A})}{\int_0^1 P(no\ detections|q_{H_A})\, dq_{H_A}}$$

We can further simplify this equation based on our assumption that the apparent prevalence of host infection is very low $(q_{H_A} << 1)$.

The equation for $P(no\ detections|q_{H_A})$ can be written as:

$$P(no\ detections|q_{H_A})$$
$$= (1-\theta_{pH})^{N_H}(1-\ \theta_{pV})^{N_V}\prod_{i=1}^{K}\left(1-\left(1-\left(\frac{\theta_{nH}}{1-\ \theta_{pH}}\right)\right)q_{H_A}Z_i\right)^{N_H}$$
$$\cdot\left(1-\left(1-\left(\frac{\theta_{nV}}{1-\ \theta_{pV}}\right)\right)\left(1-(1-\varphi_A q_{H_A}Z_i)^M\right)\right)^{N_V}$$

For small $q_{H_A}$ this can be approximated by:

$$P(no\ detections|q_{H_A})$$
$$= (1-\theta_{pH})^{N_H}(1-\ \theta_{pV})^{N_V}\prod_{i=1}^{K} exp\left(-\left(1-\left(\frac{\theta_{nH}}{1-\ \theta_{pH}}\right)\right)q_{H_A}Z_iN_H\right)$$
$$\cdot\, exp\left(-\left(1-\left(\frac{\theta_{nV}}{1-\ \theta_{pV}}\right)\right)\left(1-(1-\varphi_A q_{H_A}Z_i)^M\right)N_V\right)$$

Using the Bayesian approach described above, under the special case where $M=1$, we obtain an exponential probability distribution:

$$P\left(q_{H_A}|no\ det\right) = Be^{-Bq_{H_A}}$$

$$B = \left(\left(1-\left(\frac{\theta_{nH}}{1-\ \theta_{pH}}\right)\right)N_H + \left(1-\left(\frac{\theta_{nV}}{1-\ \theta_{pV}}\right)\right)\varphi_A N_V\right)\sum_{i=1}^{K} Z_i$$

From this, we can estimate the $x\%$ confidence interval of the apparent prevalence $q \in \left[0, q_{H_Ax}\right]$ where $q_{H_Ax}$ is found from:

$$1-\frac{x}{100} = \int_{q_{H_Ax}}^{1} P\left(q_{H_A}|no\ det\right)\,dq$$

Which for the exponential density gives:

$$q_{H_Ax} = \frac{-\ln\left(1-\left({}^{x}/_{100}\right)\right)}{B}$$

$q_{H_Ax}$ can be interpreted as the "maximal" expected apparent host prevalence when no detections are made during the monitoring program.

### Estimating the true prevalence

Our methods so far have considered the "apparent prevalence" – that is, the proportion of detectable hosts and vectors – since we are considering the ability to detect infection. However, the maximal expected apparent host prevalence may not be particularly useful for interpretation, since there is generally a period between infection and being able to detect infection in plant hosts. We term this period the "detection lag", which will be equal to the presymptomatic period before symptoms develop when we consider visual inspection as our detection strategy. A more useful measure would be the maximal true prevalence in hosts, $q_{H_Tx}$. In order to estimate this, we need to incorporate the detection lag period in the probability density $P\left(q_{H_A}|no\ det\right)$. This amounts to a transformation of variables. Considering a detection lag of $\sigma_H$, although we assume that the numbers of detectable infected hosts is low and that $q_{H_A}$ is still in the early exponential phase of growth, this may not be the case for $q_{H_T}$.

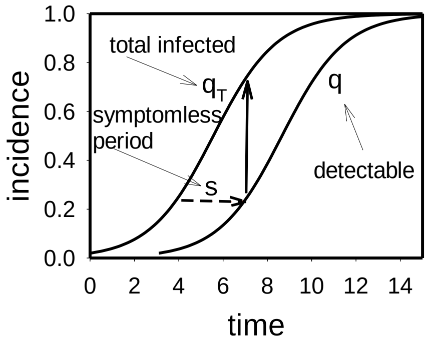


We therefore assume that $q_{H_T}$ is growing logistically. In that case the relationship between the true and apparent prevalences is given by:

$$q_{H_T} = \frac{q_{H_A}\, e^{r\sigma_H}}{1 + q_{H_A}(e^{r\sigma_H} - 1)}$$

Using this relationship, we can calculate the true host prevalence in a monitoring program where no infection has been detected in either hosts or vectors.

For the case where we only consider the first sampling point, $K = 1$ with $Se = 1 - \theta_n$ and a perfect test specificity $(\theta_p = 0)$:

$$q_{A_H} = \left(\frac{-\ln\left(1 - {}^{x}/_{100}\right)}{\left(Se_H N_H + Se_V N_V \left(\frac{D_H \beta_V}{g_0}\right)\right)}\right)$$

Using the equation above, we can estimate $q_{H_T}$ as follows:

$$q_{T_H} = \left(\frac{-\ln\left(1 - {}^{x}/_{100}\right) e^{r\delta}}{\ln\left(1 - {}^{x}/_{100}\right)(1 - e^{r\delta}) + \left(\left(\frac{D_H \beta_V}{g_0}\right) Se_V N_V + Se_H N_H\right)}\right)$$

From this, we can estimate the required number of samples from hosts and/or vectors:

$$\left(\left(\frac{D_H \beta_V}{g_0}\right) Se_V N_V + Se_H N_H\right) = -\ln\left(1 - {}^{x}/_{100}\right)\left(\left(\frac{e^{r\delta}}{q_{T_H}}\right) + (1 - e^{r\delta})\right)$$

$$N_H = \left(\frac{-\ln\left(1 - {}^{x}/_{100}\right)}{Se_H}\right)\left(\left(\frac{e^{r\delta}}{q_{T_H}}\right) + (1 - e^{r\delta})\right) - \left(\frac{Se_V}{Se_H}\right)\left(\frac{D_H \beta_V}{g_0}\right) N_V$$

$$N_V = \left(\frac{-\ln\left(1 - {}^{x}/_{100}\right)}{\left(\frac{D_H \beta_V}{g_0}\right) Se_V}\right)\left(\left(\frac{e^{r\delta}}{q_{T_H}}\right) + (1 - e^{r\delta})\right) - \left(\frac{Se_H}{Se_V}\right)\left(\frac{g_0}{D_H \beta_V}\right) N_H$$

The total sampling cost $(C)$ is the product of the sample size $(N)$ and the per-host cost $(c)$:

$$C = Nc$$

Therefore, the total cost of host-only sampling can be estimated as:

$$C_H = \left(\frac{-\ln\left(1 - {}^{x}\!/_{100}\right)}{Se_H}\right)\left(\left(\frac{e^{r\delta}}{q_{T_H}}\right) + \left(1 - e^{r\delta}\right)\right)c_H$$

And that for vector-only sampling as:

$$C_V = \left(\frac{-\ln\left(1 - {}^{x}\!/_{100}\right)}{\left(\frac{D_H\beta_V}{g_0}\right)Se_V}\right)\left(\left(\frac{e^{r\delta}}{q_{T_H}}\right) + \left(1 - e^{r\delta}\right)\right)c_V$$

Pulling these together, the ratio of the total cost of host or vector testing in order to detect by a true host prevalence of $q_{T_H}$ (assuming either hosts or vectors are sampled) is:

$$\frac{C_H}{C_V} = \left(\frac{D_H\beta_V}{g_0}\right)\left(\frac{Se_V}{Se_H}\right)\left(\frac{c_H}{c_V}\right)$$

This is not affected by the confidence level or the value of $q_{T_H}$.
