## Supplementary Tables for "Out of sight: Surveillance strategies for emerging vectored plant pathogens"

*Supplementary Table 1. Surveillance parameters*

| Parameter | Description | Value | Units | Source |
|---|---|---|---|---|
| Maximum acceptable host prevalence | | 0.01 | None | (Commission Implementing Regulation (EU) 2020/1201 of 14 August 2020 as Regards Measures to Prevent the Introduction into and the Spread within the Union of *Xylella Fastidiosa* (Wells et Al.), 2021) |
| Confidence level | 1 - probability that maximum acceptable host prevalence is exceeded | 0.90 | None | (Commission Implementing Regulation (EU) 2020/1201 of 14 August 2020 as Regards Measures to Prevent the Introduction into and the Spread within the Union of *Xylella Fastidiosa* (Wells et Al.), 2021) |

*Supplementary Table 2. Host parameters*

| Parameter | Description | Value | Units | Source |
| --- | --- | --- | --- | --- |
| Epidemic growth rate | Initial rate of exponential growth in the prevalence of infection in host | 0.0122 | New infections per infection per day | |
| Maximum acceptable host prevalence | | 0.001 | None | |
| Confidence level | 1 - probability that maximum acceptable host prevalence is exceeded | 0 | None | |
| Presymptomatic period | | 313 | Days | |
| Host laboratory test detection lag | | [varied] | Days | |
| Sensitivity of visual detection | Probability of correctly identifying an infected host visually after the asymptomatic period | 1.0 | None | |
| Sensitivity of host laboratory diagnostic | Probability of correctly identifying an infected host using a laboratory test after the detection lag period | [varied] | None | |
| Cost of visual inspection with no testing | | 2.13 | Euros per tree | Assuming that a team of two inspectors (each paid €80 per day) can inspect around 75 trees per day. Estimates from expert opinion. |
| Cost of host laboratory diagnostic | | 12.50 | Euros per test | From expert opinion (Maria Saponari). |
| Total cost of host laboratory testing | | 14.63 | Euros per test | The sum of the cost of visiting a tree and the cost |

| | | | | |
|---|---|---|---|---|
| | | | | of a laboratory diagnostic |
| Total cost of visual inspection surveillance | | 5.48 | Euros per tree | Based on estimates of proportions of (uninfected) trees which underwent ELISA testing in 2017. |

*Supplementary Table 3. Host-vector model parameters*

| Parameter | Description | Value | Units | Source |
|---|---|---|---|---|
| Prevalence in vectors | Prevalence of infection in hosts and vectors early in epidemic, measured at the same time | 0.48 | None | (Ben Moussa et al., 2016; Cornara, Cavalieri, et al., 2017; Cornara, Saponari, et al., 2017) |
| Prevalence in hosts | | 0.23 | None | Surveillance data, restricted to study areas in (Ben Moussa et al., 2016; Cornara, Cavalieri, et al., 2017; Cornara, Saponari, et al., 2017) |
| Acquisition rate ($\beta$) | Rate of vector acquisition of *X. fastidiosa* from infected hosts | 21.4428 | New vector acquisitions per vector per host per day | Fitted to data (Ben Moussa et al., 2016; Cornara, Cavalieri, et al., 2017), and an estimate of symptomatic host density. |
| Inoculation rate ($\alpha$) | Rate of host infection with *X. fastidiosa* from infected vectors | 8.579169e-06 | New host inoculations per vector per host per day | |
| Host density ($D$) | Density of hosts in an orchard/grove | 1/81 | Hosts per m$^2$ | |
| Maximum vector density | Peak vector density | 20 | Vectors per m$^2$ | (Di Serio et al., 2019; EFSA Panel on Plant Health et al., 2019) |
| Initial relative density at time of adult emergence | Initial density of vectors | 0.005343 | No units | |
| Rate of emergence of adult vectors at time of adult emergence ($g_0$) | | 0.064913 | No units | Fitted to data (Ben Moussa et al., 2016; Cornara, |

| | | | | |
|---|---|---|---|---|
| | | | | Cavalieri, et al., 2017) |
| Rate of mortality of adult vectors $(a)$ | | 0.0004574 | | Fitted to data (Ben Moussa et al., 2016; Cornara, Cavalieri, et al., 2017) |
| Time of adult emergence $(t_{em})$ | | 80 | Julian day | |
| Time of first adult infection $(t_{in})$ | | 150 | Julian day | |
| Vector laboratory test detection lag | | 0 | Days | |
| Sensitivity of vector diagnostic | Probability of correctly identifying an infected vector using a laboratory test after the detection lag period | 0.82 | No units | Taken from data on vector PCR testing (Poliakoff, 2019; Saponari et al., 2019) |
| Cost of collecting vectors | Cost of collecting a single vector | 0.76 | Euros per vector | Assuming that an inspector (paid €80 per day) can collect around 105 insects per day. Estimates from expert opinion. |
| Cost of testing vectors | Cost of the qPCR test for vector infection | 27.50 | Euros per test | Estimated cost of qPCR test. Estimates from expert opinion. |
| Vector pool size | Number of vectors pooled per test | 5 | Vectors | Estimates of maximum pool size for qPCR from expert opinion. |
